# Augmenting Flexibility: Mutual Inhibition Between Inhibitory Neurons Expands Functional Diversity

**DOI:** 10.1101/2020.11.08.371179

**Authors:** Belle Liu, Alexander James White, Chung-Chuan Lo

## Abstract

Rapid, flexible response to an ever-changing environment is critical for an organism’s survival. Recently, multicellular recordings have shown that this rapid, flexible switching between activity patterns is present in neural microcircuits. However, the underlying neural mechanism is not clear. Strikingly, we show in a neural circuit model that mutually inhibitory connections are crucial for rapid and flexible switching between distinct functions without synaptic plasticity. Here, we develop a theoretical framework to explain how inhibitory recurrent circuits give rise to this flexibility and show that mutual inhibition doubles the number of cusp bifurcations in small neural circuits. As a concrete example, we study a commonly observed class of functional motifs we call Coupled Recurrent Inhibitory and Recurrent Excitatory Loops (CRIRELs). These CRIRELs have the advantage of being both multifunctional and controllable, performing a plethora of unique functions. Finally, we demonstrate how mutual inhibition maximizes storage capacity for larger networks.

## Introduction

All organisms need to flexibly and rapidly respond to their environment. This is best accomplished by having a single network perform multiple functions (*1–13*) without needing to rely on synaptic plasticity to rapidly switch between functions (*1, 4, 12, 14–17*). It has been hypothesized that a key component of this flexibility is the recurrent nature of neural connections (*1, 2, 6, 8, 9, 14, 17–22*). Often, the focus of recurrent structures is on mutual excitation (*6, 20, 23–26*) or feedback inhibition (*6, 24, 27–33*), whereas yhe role of mutual inhibition tends to receive less attention (the term “recurrent” has been used interchangeably across literature – see Fig. 1a for how it is defined in this study), with past computational studies often focusing on slower mutual inhibitions role in central pattern generation (*12,13,34*). In addition, an accumulation of evidence has shown the preponderance of connections between inhibitory interneurons in real brain networks (*27,35–38*). These mutual inhibitory connections are found across species, from mammals (*27, 35, 37*) to simple organisms like flies (*38*). However, recent advances in multicellular recordings (*35, 39–41*) and simulation studies (*19, 39*), has given rise to new evidence that suggests that faster mutual inhibition between inhibitory inter-neuron plays a more crucial role than previously thought.

**Fig. 1.**
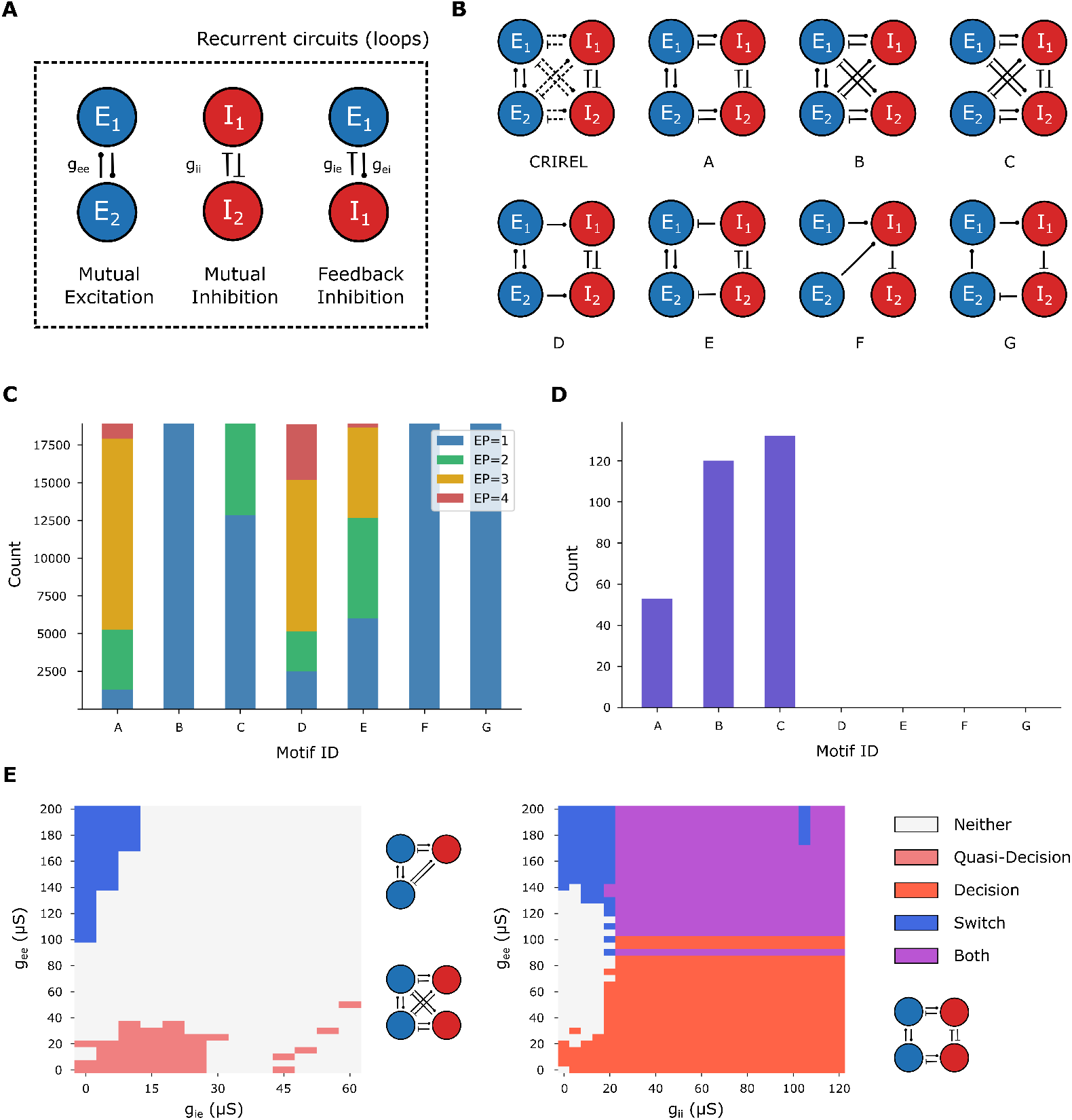
Motifs and statistical analysis of dynamical diversity. (a) Definition of the terms “recurrent” circuits, “mutual” excitation/ inhibition and “feedback” inhibition used in this paper. Synaptic weights used in part of the results are assigned based on its pre- and post-synaptic neurons, hence there are four types of weights: excitatory-to-excitatory (*g_ee_*), excitatory-to-inhibitory (*g_ei_*), inhibitory-to-inhibitory (*g_ii_*) and inhibitory-to-excitatory (*g_ie_*). (b) The motif labeled “CRIREL” is the definition of CRIREL circuits. The rest of the motifs are indexed by alphabets in no particular order. Blue represents excitation neurons, and red inhibition. (c) Parameter sweep for equilibrium points in different motifs across some parameter space. In this graph, 1 equilibrium point is in blue (EP=1), 2 is green (EP=2), 3 is yellow (EP=3) and 4 is green (EP=4). (d) Parameter sweep for CPGs in different motifs across some parameter space. (e) Parameter sweep for motifs A and B. For motif B (left), the relevant parameters concerning quasi-decision making and switch are *g_ee_* and *g_ie_*. For motif A (right), the relevant parameters are *g_ee_* and *g_ii_*. Blue represents regions where the circuit can perform switching, coral-red represents quasi-decision making, red represents decision making, and white represents no functions present. The purple region is the regime in which the system can perform both functions simultaneously.

It has been widely reported that inhibitory neurons increase the variability of a network (*3, 19, 27,42–45*), and supporting evidence from connectome studies revealed that mutual inhibition is abundant and formed locally (*35, 38, 41, 42*), possibly performing local computations (*27*). Furthermore, evidence shows that there are functional differences between feedback and mutual inhibition. For example, feedback inhibition is composed of an interconnected pair of excitatory and inhibitory neurons, which balances the network (*30–32, 46*), allowing it to approximate an arbitrary motor sequence (*14, 33*) and are useful in gain control. On the other hand, recurrent inhibition (either feedback or mutual inhibition) increases the number of basins of attraction (*6*), allowing the network to perform functions such as winner-take-all decision (*6,47–50*), bistable perception (*39,40*), oscillations (*21,22,51*), associative memory (*19,52*) and grid formation (*53*). This result, along with supporting connectome studies, all point to the idea that mutual inhibition is crucial in expanding a network’s functional repertoire, by increasing and manipulating the number of basins of attraction.

Inspired by the recent studies on mutual inhibition, we set out to systematically model its effect on a network’s functionality and dynamics. To illustrate this argument, we investigated the dynamics of smallest circuit that contains both mutual excitation and mutual inhibition. We then examine these 4-neuron circuits with and without recurrent mutual inhibition present in the network. Strikingly, by including mutual inhibition, even a small 4-neuron circuit is capable of rapidly and flexibly switching between a wide range of computations. Thus, we study a family of recurrent circuits with both a mutual inhibitory connection and a mutual excitation connection. We dubbed such 4-neuron circuits CRIREL (Coupled Recurrent Inhibition and Recurrent Excitation Loop) circuits.

We show a wide variety of computations can be performed by introducing mutual inhibition into these small 4-neuron CRIREL circuits. In particular, it can introduce new functions while subsuming the functions originally present in the circuit without mutual inhibition. We hypothesize that mutual inhibition can increase the flexibility of a 4-neuron circuit by increasing the number of bifurcations the network is near. To this end, we use phase plane analysis to understand the mechanisms that allow flexible transitions between functions, and classify different functionalities. Finally, we extend our study to large random networks and show that mutual inhibition is critical for increasing the entropy of its working memory state, implying that mutual inhibition diversifies the dynamical landscape of networks.

## Results

### Effects of recurrent structures on the complexity of circuit dynamics

We compared several circuit architectures to investigate how different recurrent structures contribute to a circuit’s dynamics (Fig. 1a,b). The first metric we employed to characterize the diversity of circuit behavior was the number of equilibrium points. If the circuit contains multiple equilibrium points, it is more likely to lead to complex activity patterns in response to different inputs, an indication of the circuit’s computational ability.

We begin with the CRIREL circuit, which contains all three recurrent structures, then remove these connections one at a time. Motif A contains all the recurrent structures contained in the other motifs (Fig. 1a). Motif B does not contain mutual inhibition, and motif C does not have mutual excitation. In motifs D and E, the feedback inhibition loop is broken – in motif D, feedback inhibition is removed, and in motif E, feedback excitation is removed. Motif F is a pure feedforward structure that was chosen as arbitrarily, and motif G is a feedforward structure that forms a single loop.

For each of the motifs, we performed a parameter sweep over synaptic weights and bias currents. We ran a total of 18900 trials. The protocol detects equilibrium points by stochastically exploring the system in phase space. The firing rate sequence of the trial is then put though the Elbow method to judge the number of clusters that are formed. These clusters correlate to the amount of stable equilibrium points in the system.

The results show that motif A has the largest parameter regime in which the system not monostable, i.e. the number of equilibrium points exceed 1 (Fig. 1c; monostable in blue). Furthermore, motifs that have mutual inhibition (motifs A, C, D, E) have parameter sets with equilibrium point counts larger than two, while those with mutual excitation but not mutual inhibition has at most counts of two, and feedforward motifs have no bistability at all. In contrast, the presence or absence of feedback inhibition does not make such a large difference. For example, motifs A, D and E all have high numbers of equilibrium points, but D and E do not contain feedback inhibition while A does.

While the first metric shows the number of equilibrium points in the network, it does not reveal whether other stable structures are possible within the network, such as oscillation given constant input. Therefore, the second metric we employed tested whether central pattern generators (CPG) are present in the network. We count the number of distinct inter-spike intervals (ISI) within the inhibitory neurons, and if the number of ISIs is greater than 1, that implies that there is a CPG (see methods for more details). This analysis shows that CPGs are only found in A, B and C – the ones that contain feedback inhibition (Fig. 1d).

In summary, we illustrated the functional differences between mutual and feedback inhibition – mutual inhibition increases the number of equilibrium points in the network, while feedback inhibition allows the network to oscillate when given constant input.

### Functional differences between feedback and mutual inhibition

The statistical results indicate that mutual inhibition increases the dynamical states, but does not explain the mechanism behind it. Hence here we consider an explicit example that compares feedback inhibition with mutual inhibition, and compare motif A with motif B. Specifically, we compared the two motifs’ ability to perform different functions within some relevant parameter regime.

Motif B contains mutual excitation, which allows it to perform a switch-like function, where some brief positive current pulse can transfer the system to the ON state, and a negative pulse can turn it off. Motif B also has feedback inhibition, which skews the basin of attraction one way, therefore making one of the excitatory neuron fire faster than the other, performing a quasi-decision like function. However, these two functions cannot coexist within the circuit (Fig. 1e, left panel). If we add recurrent inhibition to the circuit, however, this problem can be resolved. Consider motif A, which has mutual excitation, feedback inhibition, as well as mutual inhibition. This gives a large parameter regime in which motif A contains both functions simultaneously (Fig. 1e, right panel). Here, we stress that even though motif A has more synapses, it is motif B (with mutual inhibition) that is more functional. Therefore, presence of mutual inhibition can add more functions to the underlying neural network.

### Dynamics of CRIREL circuits

All of the statistical analysis above point to mutual inhibition being crucial in expanding the functional repertoire of a network. However, to have a better grasp on the functions CRIREL circuits actually bring about, we must look into its dynamics. To this end, we reduced the 4-dimensional CRIREL circuit into a 2-dimensional reduced model, with each of the recurrent circuits compressed into 1 dimension (see Methods and Supplementary Note). Conceptually, it can be understood as preserving the dimensions that are important to the dynamics of interest by considering the properties of the two mutual connections. Because each LIF neuron as a class 1 excitable neuron (*54*), we can approximate each neuron as a Wilson-Cowen (*21, 55*) type firing rate model, with each neuron’s firing rate as *e*_1_, *e*_2_, *i*_1_ and *i*_2_. Since mutual excitation tends to synchronize, the dimension of *e*_1_ − *e*_2_ is less important than *e*_1_ + *e*_2_ if the coupling is much weaker than the mutual connection strength, where *e_i_* represents the firing rate of excitatory neuron *i*, which is denoted as *E_i_*. On the other hand, mutual inhibition desynchronizes, therefore *i*_1_ + *i*_2_ is less important than *i*_1_ − *i*_2_ when the coupling is weak. Similarly, *i_j_* represents the firing rate of inhibitory neuron *j*, which is denoted as *I_j_*. Hence, by retaining the dimensions of *e*_1_+*e*_2_ and *i*_1_ − *i*_2_ only, we can gain a geometrical understanding of how the system operates, which in turn sheds light on the potential computations recurrent circuits can perform (Fig. 2a).

**Fig. 2.**
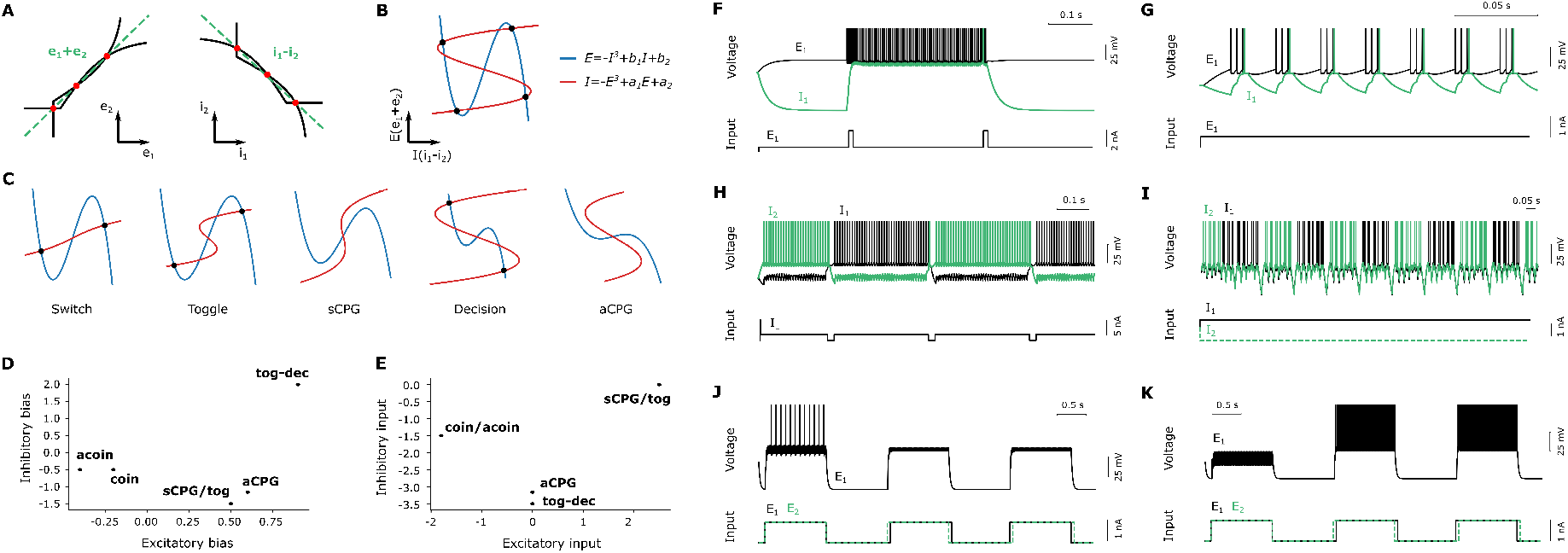
The reduced model of the CRIREL circuit. (a) The phase diagrams of the mutual excitation (left) and inhibition loop (right) when decoupled from one another. The black curves are nullclines and the spots represent equilibrium points. The green dashed line shows the direction in which in interesting dynamics lie on (it is only approximate for the inhibitory case). (b) When weakly coupled together, the CRIREL circuit can be reduced to two dimensions. The blue curve is the nullcline of the mutual excitation loop, red is the nullcline of mutual inhibition, and black dots are stable equilibria. (c) By adjusting the four parameters *a*_1_, *a*_2_, *b*_1_ and *b*_2_, the reduced model will change into different configurations, each of them corresponding to different functions. Demonstration of rapid flexibility using bias currents. All functions coexist for a single set of synaptic weights, namely *g_ee_* = *g_ii_* = *g_ei_* = *g_ie_* = 70 *mS*. (d) The plot of excitatory and inhibitory bias currents. Each Dot shows the location in bias current parameter space for each function. (e) The plot of input magnitude excitatory and inhibitory for each function. (f) Demonstration of the toggle function using square pulses into the excitatory subsystem. (g) Demonstration of the synchronized CPG function, note that there is a no input, only bias current. (h) Demonstration of the toggle-decide function using pulses into the inhibitory subsystem. (i) Demonstration of the anti-synchronized CPG, which again has no input into the network. (j) Coincidence detection with input into both excitatory and inhibitory subsystems. (k) Anticoincidence detection using input into both subsystems.

The resulting reduced model (Fig. 2b) can be visualized using the coordinates **E** ≈ *e*_1_+*e*_2_ and **I** ≈ *i*_1_ − *i*_2_. The blue and red line represents the nullclines of **E** and **I** respectively. Note that both nullclines are of a cubic form, which is the normal form of a cusp bifurcation, thereby proving that mutual connections do indeed add cusp bifurcations to the circuit. Since there are two cubic functions, we know that this system has a double-cusp bifurcation. There are four free parameters in this reduced model, which comes from complicated nonlinear mappings of the original neuronal and bias current parameters. For physical intuition, *a*_1_ and *b*_1_ are nonlinear mappings of structural parameters of the neurons, such as capacitance or time constant or synaptic weights. *a*_2_ and *b*_2_ are mappings from the bias currents into the circuit. These parameters allow the cubic functions to change shape as well as translation, which gives rise to many different phase diagrams (Fig. 2c). These phase diagrams correspond to different “modes” of the circuit, in which it performs different functions. We will go over all of the functions we discovered below, but note that this is by no means a complete list of what the circuit is capable of doing.

### Rapid flexibility using bias currents to control functionality

Next, we consider a network flexible if multiple functions coexist for a given set of synaptic weights. We are able to reproduce all functions listed in (Fig. 2d-e) with a single network with a single set of synaptic weights. For demonstration purposes, we chose the synaptic weights *g_ee_* = *g_ii_* = *g_ei_* = *g_ii_* = 70 *mS*.

Strikingly, we are able to use bias current alone to switch between various functions (Fig. 2 2c). Importantly, this endows our network with a set of 6 unique functions (Fig. 2f-j) that can easily and flexibly switched between by modulating bias current and input type. The six functions are toggling (Fig. 2f), synchronized CPG (Fig. 2g), toggle-decide (Fig. 2h), anti-synchronized CPG (Fig. 2i), coincidence detection (Fig. 2j), and anti-coincidence detection (Fig. 2k).

A nice consequence of this rapid flexibility is to change the functionality on a timescale corresponding to the membrane time constant (here 20 *msec*). It’s worth stressing that this does not require a slower change in synaptic weight, and that every function is easily accessible from the other functions.

The functions of CRIREL circuits include rudimentary ones that rely more heavily on mutual excitation, such as switch, toggle and synchronized CPG; it also includes ones in which computations are mainly performed in the mutual inhibition loop, such as decision-making and anti-synchronized CPG. Building from these concepts, we can also have computationally interesting functions, such as working memory, threshold-based filtering and timing-based decision-making.

#### Computations in the excitatory subsystem

##### Switch

We first demonstrate one of the simplest computations, which is a switch-like function (Fig. 3a). An intuitive understanding of switching and its behavior is already described in previous sections. Here we demonstrate how to comprehend this function geometrically using the reduced model. The reduced model for switch has only two stable equilibrium points separated by a saddle equilibrium (Fig. 3b). The simulation begins at the equilibrium point where all the neurons are silent, i.e. the OFF state. When a brief positive current pulse is given to one or both of the excitatory neurons, the **E**-nullcline shifts upwards, annihilating the OFF state. Therefore the system transits to the remaining equilibrium point, i.e. the ON state, and remains there even as the input is removed and the **E**-nullcline returns to its original position. This means that the circuit will continue firing even in the absence of input. When a negative current pulse is applied, the opposite happens, and the system returns to the OFF state. This shows that the circuit is bistable. We see that the reduced model is consistent with the intuitive explanation given above (Fig. 3b). Moreover, we tested that this switch into the upstate was present for a broad range of potential input stimuli (Fig. 3c).

**Fig. 3.**
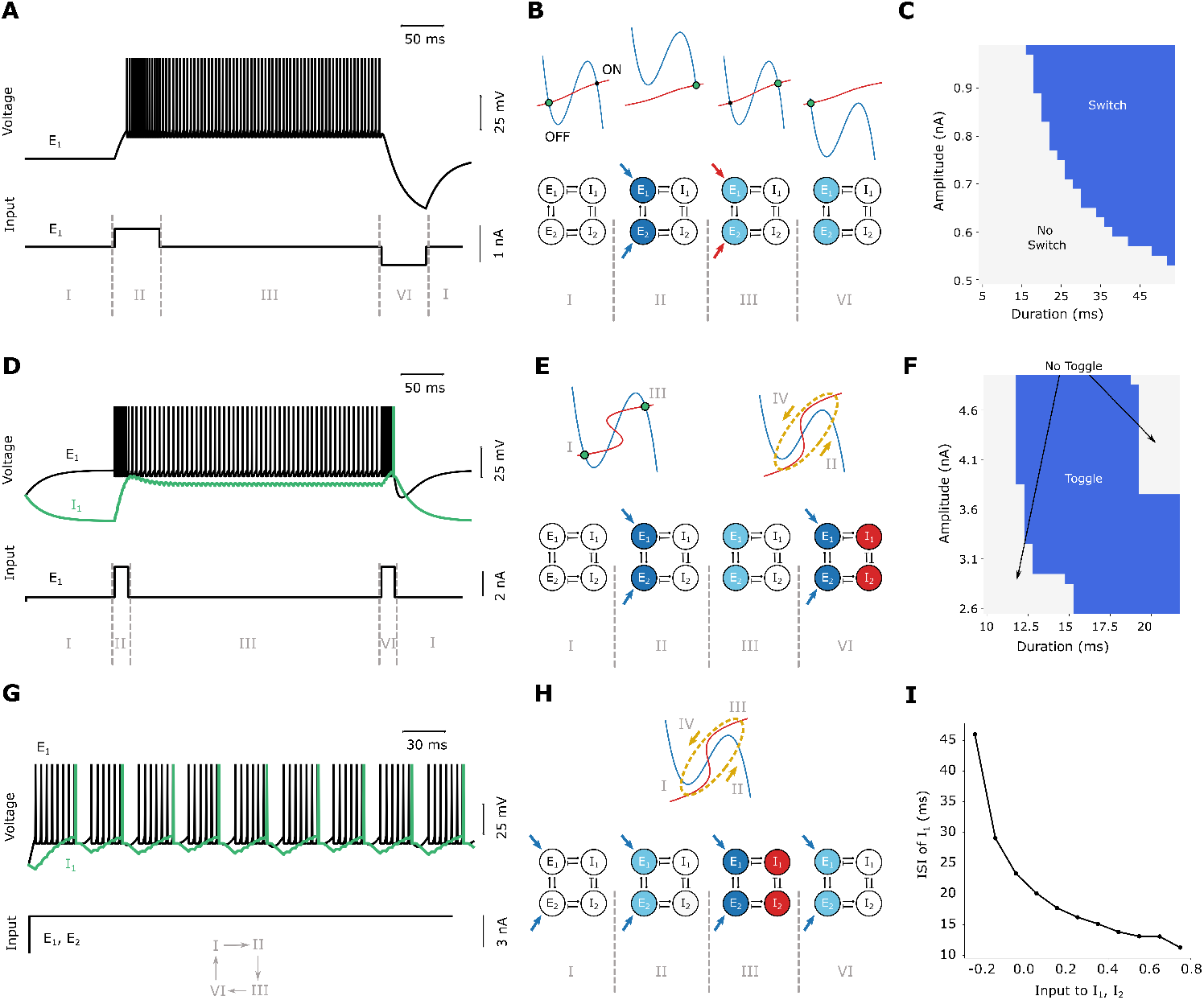
Switch, toggle and synchronous CPG. (a) Switch voltage traces. Top row: the voltage trace of neuron *E*_1_. Bottom row: the current input into *E*_1_. The roman numerals represent different phases of the input, and corresponds to the roman numerals shown in panel (b). (b) Switch phase diagrams and schematics of neural activity. Top row: switch function phase diagram. The green dot represents the state of the system. Bottom row: Schematic diagram of the relative activity of each neuron during different time periods. Blue represents excitation, and red inhibition. The darker the color, the higher the activity. (c) Inputs that induce a switch. A parameter sweep over duration and amplitude of the input pulse that induces a switch into the upstate. Blue represent stimuli that induce a switch, while gray represents those that do not. (d) Toggle voltage traces. Top row: the voltage traces of *E*_1_ (in black) and *I*_1_ (in green). Bottom row: the current given to *E*_1_. The roman numerals correspond to the roman numerals shown in panel (e). (e) Toggle phase diagrams and schematics of neural activity. Top row: toggle function phase diagram. Bottom row: Schematic diagram of the firing rates of each neuron during different time periods. (f) Inputs that induce a toggle. A parameter sweep over duration and amplitude of the input pulse that toggles between the upstate and downstate. Blue represent stimuli that induce a switch, while gray represents those that do not. (g) synchronous CPG voltage traces. The rows are identical to those of (d). Here, however, the current into E1 is changed from pulses into a constant input. The roman numerals arranged in a loop means that the four phases are occurring repeatedly. (h) sCPG phase diagrams and schematics of neural activity. (i) ISI of inhibitory spikes. Parameter sweep over the inhibitory bias current, and its effect on the period of the CPG.

#### Computations in the excitatory subsystem with feedback inhibition

##### Toggle

A function that is similar to switch is toggle, except that it can be turned off by a positive current pulse instead of a negative pulse. We can visualize this by plotting the voltage traces of one of the excitatory neurons (which are synchronized) and one of the inhibitory neurons (Fig. 3d). The first pulse causes the system to go to the ON state, and the firing rate of the inhibitory neurons are insufficient to inhibit the excitatory loop. However, when a second identical pulse enters the system, the firing rate of the excitatory loop increases, and concurrently the inhibitory loop. This extra push is enough for the inhibitory neurons to turn off the excitatory neurons, thus returning the whole system to inactivity.

Again, we can gain insight by examining the phase portrait of the system. The simple shifting mechanism of the switch-system no longer applies here. As the impulse current excites the excitatory loop and consequently the inhibition loop, the inhibitory response deepens, causing the inhibitory nullcline to change its shape (Fig. 3e). This in turns causes both the ON and OFF state to annihilate, and the system goes from 5 equilibrium points to one unstable spiral node. This means that for a brief amount of time, there exists a limit cycle and the system’s state rotates. When the impulse input ends, the inhibitory system relaxes and the 5 equilibrium points return, but by that time the system is already trapped in the ON state. The second pulse repeats the process and rotates the system yet again, allowing it to return to the OFF state. We also examine the required amplitude and duration of stimuli required to toggle the system (Fig. 3f). Interestingly, for very long duration the system may rotate a full rotation, rather than a half rotation. This will cause the system not to toggle for long duration stimuli.

##### Synchronized Central Pattern Generator

The presence of toggle implies that the circuit is also capable of sustaining periodic oscillations, that is, capable of becoming CPGs. Note that if the impulse current is changed into a constant bias current, the system will repeat the process of toggling over and over again, thus undergoing a synchronized bursting behavior (Fig. 3g). The phase diagram of synchronous CPG is the limit-cycle mode (Fig. 3h). Moreover, we showed that inhibitory bias current can tune the network over a wide operating frequency (Fig. 3i).

#### Computations in the inhibitory subsystem

##### Decision making

Switch, toggle and synchronous CPG rely heavily on the dynamics of the mutual excitation loop. Decision making, on the other hand, is mainly due to mutual inhibition. For a decision network, when two inputs are injected into the neurons *E*_1_ and *E*_2_, the larger of the two inputs will “choose” their corresponding inhibitory neuron, thus preventing the other inhibitory neuron from firing (Fig. 4a). Logically, this is the exact counterpart of the switch system, the difference being that the nonlinearity now lies in the **I**-nullcline instead of the **E**-nullcline. While inputs shift the **E**-nullcline either up or down in the switch or toggle phase diagram, it shifts the inhibitory neuron either left or right in this case, hence forcing the system to decide between the two equilibrium points. Geometrically, this corresponds to “rotating” the nullclines of the synchronized CPG by 90 degrees (Fig. 4b). To quantify the performance of decision making, we examined the accuracy and reaction time of the circuit (Fig. 4c). The results are consistent with decision making networks containing two competing populations of excitatory neurons – as the input becomes more coherent, accuracy increases and reaction time decreases.

**Fig. 4.**
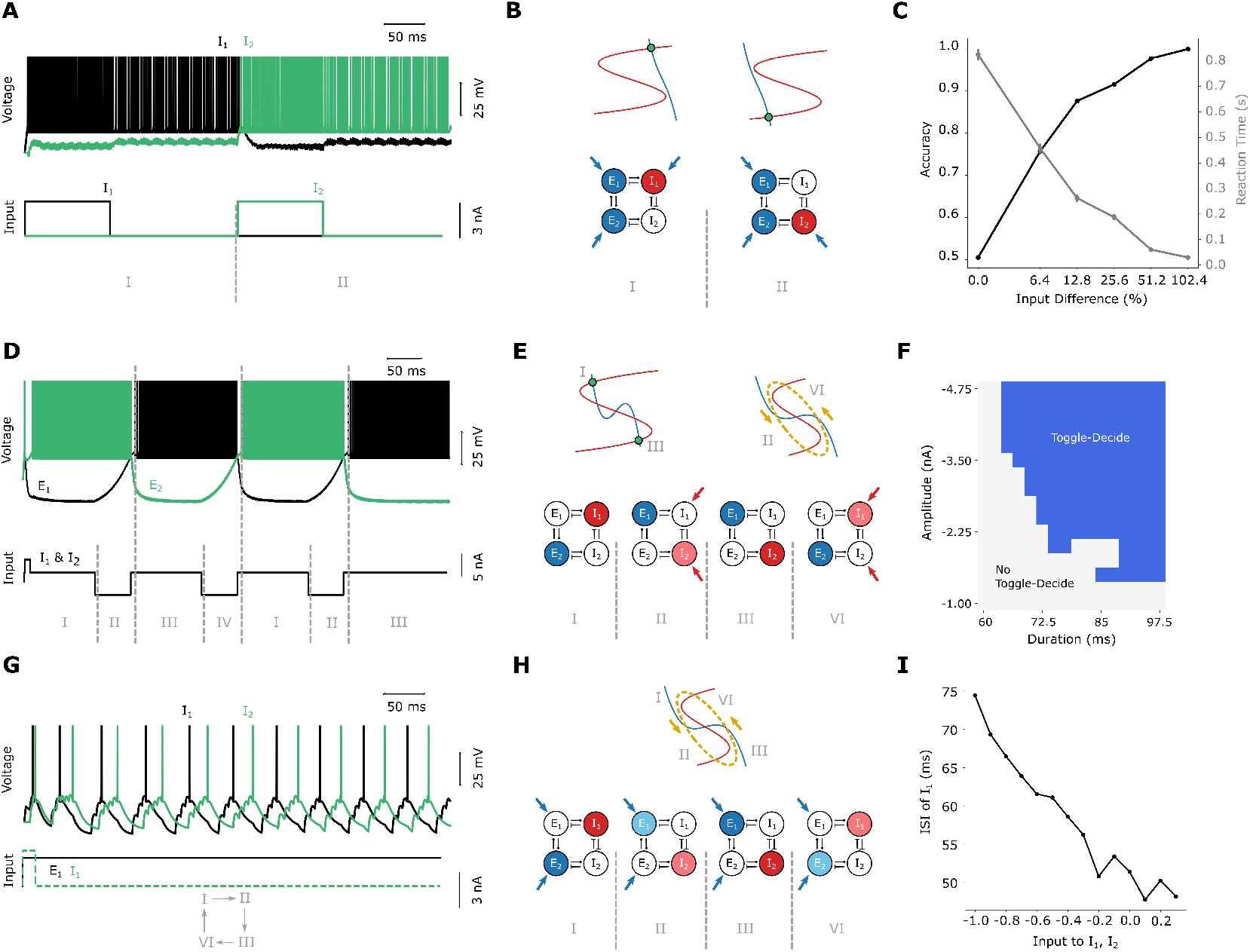
Decision making and asynchronous CPG. (a) Decision making voltage traces. Top row: the voltage traces of *I*_1_ (in black) and *I*_2_ (in green). Bottom row: the current given to *I*_1_ (in black) and *I*_2_ (in green). Here, *E*_1_ and *E*_2_ are given constant bias currents to maintain the decision after the input is removed. (b) Decision making phase diagrams and schematics of neural activity. (c) Psychometric function of decision making. As the input difference of the two inputs increases, the accuracy (black) increases and the reaction time (grey) decreases. The error bar for the reaction time represents its standard error. (d) Anti-toggle voltage traces. (e) Anti-toggle phase diagrams and schematics of neural activity. (f) A parameter sweep over duration and amplitude of the input pulse that anti-toggle. Blue represent stimuli that induce a anti-toggle, while gray represents those that do not. (g) Asynchronous CPG voltage traces. (h) Asynchronous CPG phase diagrams and schematics of neural activity. (i) ISI of inhibitory spikes. Parameter sweep over the inhibitory bias current, and its effect on the period of the CPG.

##### Anti-toggle (anti-synchronized toggle)

Just as decision making is the counterpart of the switch function, anti-toggle is the counterpart of toggle. By taking advantage of the reduced model, we can rotate the nullclines by 90 degrees to generate an anti-toggle from toggle. This is achieved by swapping the parameters of the excitatory and inhibitory systems. As with the case of toggling, in this mode the excitatory neurons of the excitatory sub-circuit can be turned on and off by positive input pulses. The difference is that now when one inhibitory neuron is turned off, the other one automatically turns on (Fig. 4d, e). This function is extremely robust, as it works in a wide range of parameters (Fig. 4f).

##### Asynchronized Central Pattern Generator

Similar to how toggle gives rise to synchronized CPGs, we can do the same for anti-toggle to an anti-synchronized CPG. by changing the current pulses into a constant bias. This change will not directly yield anti-synchronized CPGs, since a constant bias would be reinforcing the system to stay within the same equilibrium point. However, if we introduce time-scale separation, e.g. mechanisms such as slow GABA synapses, calcium dynamics, and second-messenger systems, the circuit can oscillate. We can visualize this by taking advantage of a limit cycle in the reduced system (Fig. 4g, h). Again, we examined how the inhibitory bias current controls the operating frequency of the CPG (Fig. 4i), and showed that it operates over a wide rage of frequencies.

#### Computations in the CRIREL system

##### Working memory

The situation described above complicates when the bistability of the excitatory sub-system is combined with decision making of the inhibitory sub-system. Since the ON state exists, this means that the decision can be remembered even after the input is withdrawn because of the excitatory sub-system. When receiving strong input, the excitation sub-system enters the ON state asymmetrically. One of the excitatory neurons will have higher firing rates, thus allowing its neighboring inhibitory neuron to fire and inhibit the other one. Furthermore, the feedback inhibition strength is not strong enough to turn the excitatory subsystem off, thus the system “remembers” the input (Fig. 5).

**Fig. 5.**
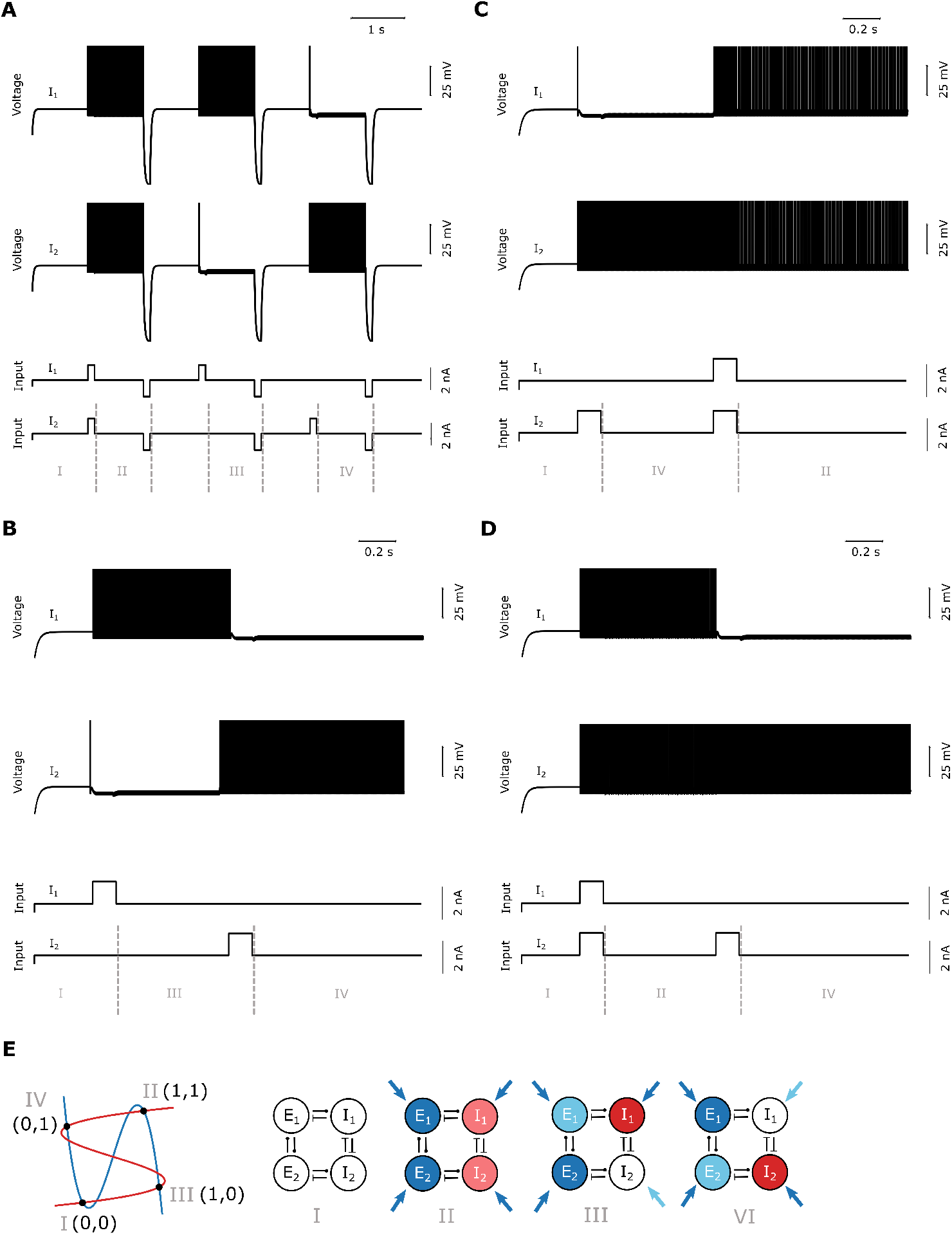
Working memory. (a) Voltage trace. This circuits stores a “two bit” signal in working memory, i.e. (1,1), (1,0), (0,1) and (0,0), corresponding to (*I*_1_ on, *I*_2_ on), (*I*_1_ on, *I*_2_ off), (*I*_1_ off, *I*_2_ on), (*I*_1_ off, *I*_2_ off). This can be seen in the top two rows of the graph, showing the voltage traces of *I*_1_ and *I*_2_. The bottom two rows are the inputs to the four neurons. (b-d) Switching memory without reset. (b) Switching from (1,0) to (0,1). (c) Switching from (0,1) to (1,1). (d) Switching from (1,1) to (0,1). (e) Working memory phase diagrams and schematics of neural activity. Top row: phase diagram. Bottom row: Schematic diagram of the neural activity.

There are two protocols to change the memory currently being maintained in the system. One can clear the system with a reset signal and the system will return to rest (Fig. 5a). The memory can also be overwritten directly (Fig. 5b-d).

The analysis of this system is identical to the decision-making system, except that there could be 4 equilibrium points (Fig. 5e). It can indeed be confirmed that this nullcline configuration is present in the full CRIREL system, where the inhibitory neurons can either be both on (i.e. (1, 1)), one on and one off (i.e. (1, 0) or (0, 1)) or both off (i.e. (0, 0)). Thus this system is capable of forming a working memory system with 2-bit memory. This is a perfect example of how simple functions such as decision making and switch can combine to yield a more interesting and useful function.

##### Threshold-based filtering

The capability of making decisions allows CRIRELs to be utilized in many different ways, including useful functions such as creating a safety net for hyperactivity. If we allow the CRIREL circuit to take on asymmetrical parameters (Fig. 6a, left) of membrane and synapse, the circuit can become a low-pass filter in terms of amplitudes, where small inputs are remembered by the circuit (working memory like), and larger ones terminate the bistability (toggle like) (Fig. 6b). This computation is possible precisely due to the presence of asymmetries in the network. When one of the excitatory neurons receive a small input, since *I*_1_ has stronger synaptic weights, *I*_1_ will be chosen and thus inhibit *I*_2_. However, when the input is sufficiently large, *I*_2_ has smaller capacitance and time constants than *I*_1_, and therefore will be excited faster than *I*_1_, thus winning the decision. The feedback inhibition from *I*_1_ is weak, therefore when *I*_1_ is chosen, it does not affect the network’s bistability; however, the feedback inhibition from *I*_2_ is strong, therefore whenever *I*_2_ is chosen, the network shuts down. Hence the circuit serves as a gateway that filters out inputs that are too large, which could be potentially useful for preventing epilepsy. The threshold of the circuit can be adjusted by manipulating the bias current of *I*_1_ (Fig. 6c).

**Fig. 6.**
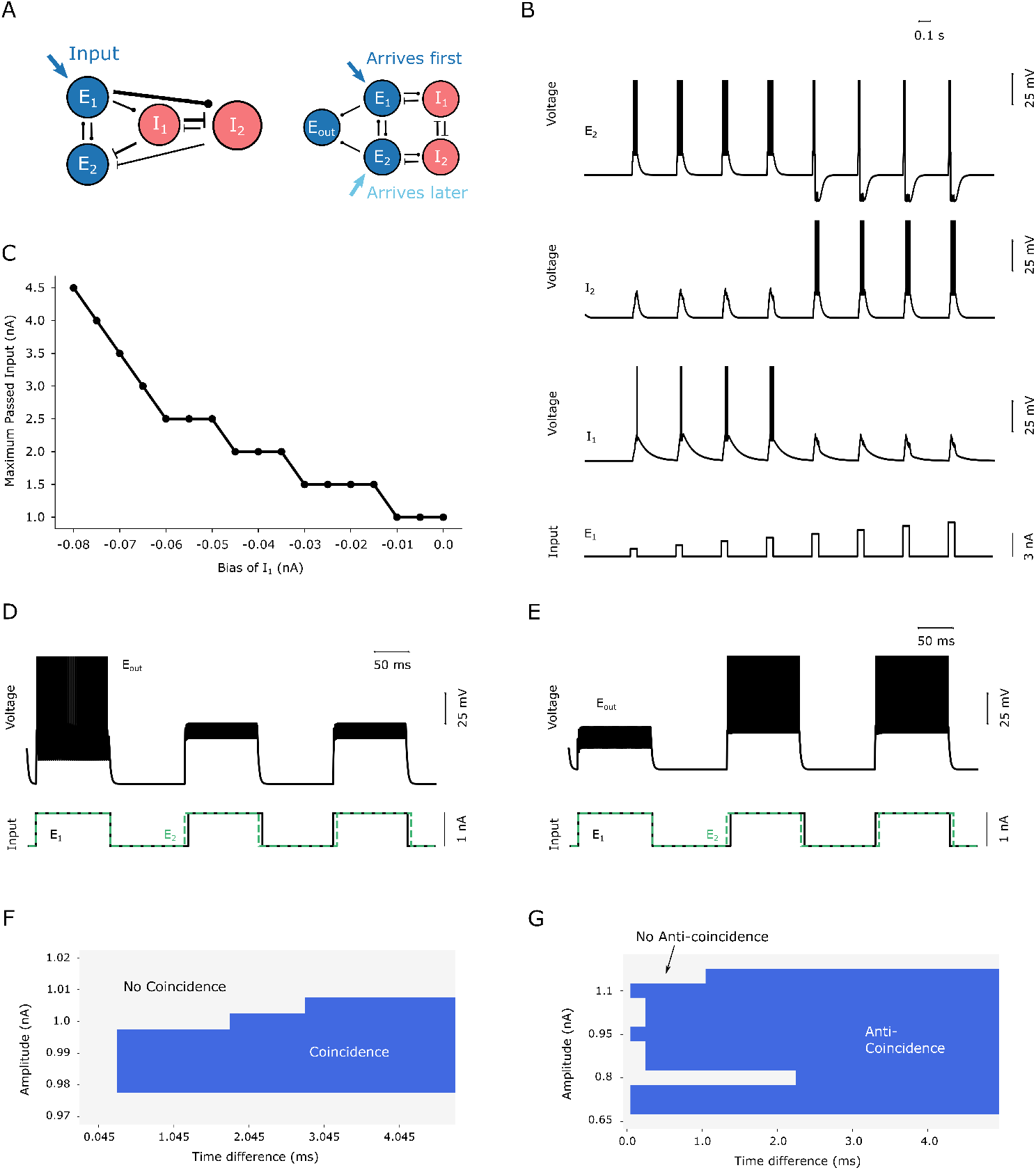
Threshold-based filtering and coincidence detection.(a) Circuit motifs for threshold-based filtering (left) and timing based detection (right). The asymmetries of the parameters are shown schematically in the diagram: thicker lines represent larger weights, and vice versa. (b) Voltage traces of threshold-based filtering. The top three row shows the voltage traces of *E*_2_, *I*_2_ and *I*_1_, and the bottom row shows the injected current into *E*_1_. Here we inject neuron *E*_1_ with different pulses of increasing amplitude. (c) Maximum amplitude of input that passes the filter as a function of bias into *I*_1_. The steps are discrete because the input we used are discrete. (d) Timing based detection, coincidence-mode. The top row is the voltage trace for the output neuron. The bottom row is the input for *E*_2_ (in black) and *E*_1_ (in green). Since the timing difference between the input is small (5 ms), for visual demonstration the difference is exaggerated. (e) Timing based detection, anti-coincidence-mode. The top row is the voltage trace for the output neuron. The bottom row is the input for *E*_2_ (in black) and *E*_1_ (in green). (f-g) A parameter sweep for both coincidence (f) and anti-coincidence (g) over both amplitude of signal and timing difference. Blue represents the existence of the function, while light gray represents its absence.

##### Timing-based detection

Aside from making decisions based on the magnitude of the input, what is perhaps even more intriguing is detecting the relative timing of inputs (Fig. 6 d,e). Even a sub millisecond difference between the inputs is sufficient for the network to perceive. Timing-based detections can be accomplished if we allow sufficiently strong feedback inhibition, as we did in toggling. Under this condition, two modes are allowed: the coincidence detection mode (Fig. 6d) and the anti-coincidence detection mode (Fig. 6e). In the anti-coincidence mode, the network will chose whichever input came first, but will turn off completely if inputs arrive at the same time. In the coincidence mode, the network will only fire if the inputs arrive simultaneously (Fig. 6e). Note that for these two modes, an additional neuron is added to the CRIREL circuit that combines the output of *E*_1_ and *E*_2_ (Fig. 6a, right). However, this extra neuron is only added for the sake of having a more clear-cut output. The computation for these two modes are still done within the CRIREL circuit itself.

Here, coincidence is controlled primarily by mutual excitation. The synaptic input is sufficient to turn the system on when both inputs arrive at the same time, but if there is a delay in the system then, the system will not fire. Conversely, the anti-coincidence system takes advantage of the toggle like dynamics, where it turns the system off if and only if both of the inputs arrive simultaneously. Otherwise, the inhibitory neurons do not fire with sufficient strength to turn the system off. Coincidence has a narrow range of operation (Fig. 6f) because it relies on a more delicate balance between the inhibitory and excitatory subsystems. One the other hand, anti-coincidence operates over a much broader range of parameters (Fig 6g).

### Memory in large network

So far, we have explicitly explained how mutual inhibition adds a cusp bifurcation to the circuit, which allows the circuit to perform various functions. Here, we extend this argument to large random networks. We focused on the function of working memory in particular, as working memory requires use of both on-off switches (excitatory cusps) as well as decision making (inhibitory cusps). We showed that an increase in mutual inhibition strength allows the network to have a wider variety of memory states. However, other functions that are present in the CRIREL circuit can also be reproduced in the large network as well (see Fig. S3).

The random network consists of 100 neurons, 75 excitatory and 25 inhibitory, where each connection has a 50 percent probability of being present (Fig. 7a). *g_ie_* and *g_ii_* are varied from 0 to approximately 30 (*μS*). For each parameter set, the excitatory neurons of the network are given random current pulses, and after the stimulation stops the network relaxes into a steady state. Since the stimulation is random, each time the steady state might be identical or different, and this is repeated for 100 times per parameter set. Each steady state is recorded as a “word”, where different words represent different steady states (Fig. 7b; see Methods). The entropy for the distribution of the words across 100 trials, *H_word_*, is calculated for the various parameter sets, which reflects how diverse the steady states are (Fig. 7c).

**Fig. 7.**
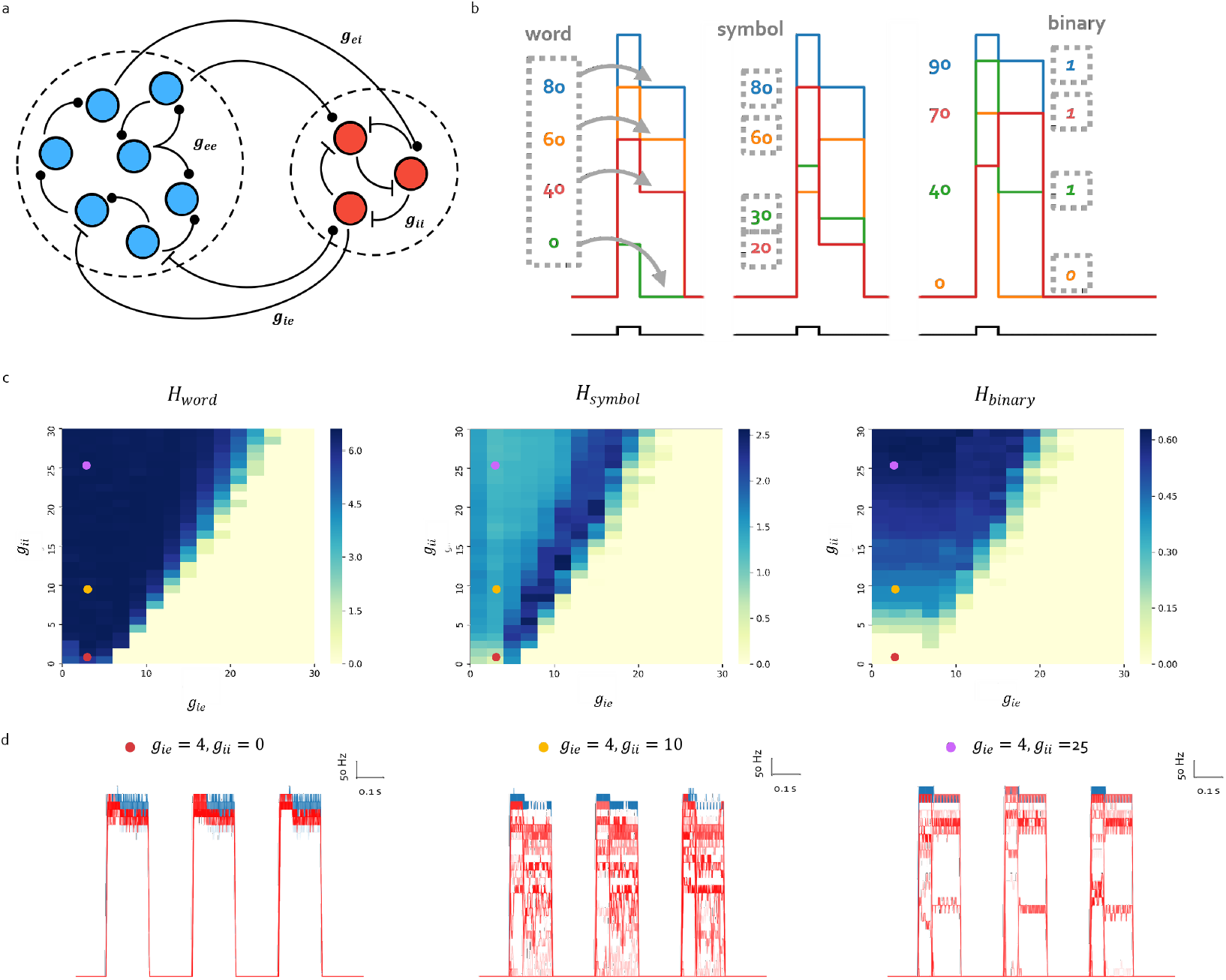
Memory in large network. (a) Schematic diagram of large random network. (b) Schematic diagram of how entropy is calculated. Each line represents the firing rate of a neuron, and the firing rate of the neuron population is defined as a “word”. The words across 100 trials forms a probability mass function, which is used to calculate *H_word_*. The word itself is also a probability mass function for different firing rates, which also has an associated entropy. The trial-average of this entropy is called *H_symbol_*. Alternatively, each neuron can be labeled as on (*1*) or off (0), and the entropy associated with this is called *H_binary_*. (c) From left to right: *H_word_*, *H_symbol_*, *H_binary_* for different *g_ii_* and *g_ie_*. (d) Firing rate traces for neurons across 3 trials, with different synaptic weights. Each trial showcases how the steady state activity differs between having different *g_ii_* values. Blue traces represent excitatory neurons, and red traces inhibitory neurons. From left to right: increasing *g_ii_*.

For the same *g_ie_* value, *H_word_* increases as *g_ii_* increases (Fig. 7c,d). To understand the reason behind this, we turn to two other entropy measures, *H_symbol_* and *H_binary_* (Fig. 7b). Simply put, *H_symbol_* measures how diverse the firing rate across population is, for a single trial. If the activities of all neurons are high (Fig. 7d left) then there are not a lot of symbols available to form a word, hence *H_symbol_* would be low and consequentially *H_word_* as well. Moreover, It can be shown that there is a regime of suitable *g_ii_* to *g_ie_* ratio where *H_symbol_* is maximized (Fig. 7c). This trend is quite different from *H_word_*, where it increases almost monotonically with *g_ii_*. Therefore, an increase in the number of symbols is not the main mechanism behind what we observed in *H_word_*.

Therefore, we used yet another method for calculating entropy, *H_binary_*. Here, if a neuron fires, it is labeled as “1”, while a silent neuron is labeled “0” (Fig. 7b). Each time the network relaxes into a steady state, it yields a distribution of 1s and 0s across the neuron population, and the entropy for that distribution is calculated. Essentially, if the neurons’ firing rates are more “all or nothing” (or “winner takes all”) – so that the ratio between the count of 1s and 0s are more equal – then *H_binary_* will be higher. Moreover, *H_binary_* is augmented as *g_ii_* increases, which follows a similar trend to *H_word_* (Fig. 7c), implying that the increase in memory capacity as shown by *H_word_* may be caused by the neurons’ firing becoming more “all or nothing”, which is in turns due to an increase in *g_ii_*.

Furthermore, we note that for large *g_ii_* values, its entropy value is near maximal. That is, if all 100 trials result in unique states, then 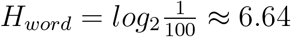, and we see that the entropy values of large *g_ii_* are near that value.

To conclude, increasing *g_ii_* increases the number of memory states the network contains. Consistent with how mutual inhibition introduces new bifurcations to the CRIREL circuit, mutual inhibition in a large network also complicates its dynamics and yields a more computationally useful product. The large network is also capable of performing other functions found in the CRIREL circuit (Fig. S3).

## Discussion

In the present study, we show that mutual inhibition plays a key role in increasing the computation complexity and functional repertoire. This is evident from: (*1*) motifs that contain mutual inhibition exhibit more functions than those without, (*2*) the ability of a specific class of the motif, called CRIREL, to rapidly and flexibly switch between multiple biologically relevant functions for the same set of synaptic weights, and (*3*) higher working memory capacity in large neural networks with mutual inhibition than those without. We also demonstrate that the key mechanism that underlying the functionality of mutual inhibition lies in its ability to increase the basins of attraction from an increase in the underlying cusp bifurcations.

This is inline with past research, as connections between inhibitory neurons has been shown in many different contexts (*27–29, 35, 41, 42, 56*). It is interesting to note that in a recent study (*38*) that reconstructed single-cell level brain-wide connectome of fruit flies (*57–59*), putative inhibitory neurons receive more inhibitory than the excitatory inputs. Moreover, studies in the hippocampus of rats have found a plethora of mutually inhibitory parvalbumin interneurons (*35, 41*). These findings imply the importance of mutual inhibition in the brain network. We have shown computationally that these types of connections are useful and perform a unique role in computational processing by expanding the number of computational states a network has. We also hypothesis the existence of a function called anti-toggle, that is capable of seamlessly switching between different basins of attraction, which is potentially useful in tracking changes of state, working memory based decisions, or even counting. Furthermore, the existence of decisions and anti-toggle in the inhibitory sub-network could potentially be one mechanism behind simultaneous suppression and excitation of inhibitory interneurons observed in various studies of cortical neurons (*27, 37, 42*).

Moreover, our results are in line with studies that have reported an increase in variability (*37, 42*) and working memory maintenance (*19*) when the connection strength of inhibition is increased. In our study, we showed that increases in *g_ii_* is more responsible for the increase in the number of steady states than *g_ie_*. This tracks with the presence of a multiple cusp bifurcation. As the large network simulations showed, for each inhibitory neuron with all-to-all inhibitory connections added, there is an additional cusp bifurcation. This doubles the number of unique combinations of states the network can perform.

At last, we note that a network’s functional capacity is not determined solely by the number of synapses (*21, 42, 46, 60*). In particular, while each type of synapse increases the amount of potential computations a network can have, the network is not necessarily flexible if it cannot seamlessly transition between different functional modes by modulating the parameters. Whether the network is capable of easy transitions is highly dependent on the underlying bifurcation, which is the phenomenon of switching between behavioral characteristics through the change of model parameters (*21, 54, 61, 62*). Specifically, our research reiterates that networks near complicated bifurcations are computationally useful (*21*), because only networks near a bifurcation have non-trivial dynamics (*14, 21,62*). In short, one can quickly and precisely control the neural circuit operations by controlling the nearby bifurcation.

One area our research can be extended to is examining the effects of gap junctions on the network. Some have found that mutual inhibition and gap junctions tend to coexist in some circuits (*35,41*). It has been shown that gap junctions have a tendency to average out the winner-take-all aspect of the decisions (*63*).

In addition, in our study we have mostly overlooked plastic synapses. Several studies have shown that this is an important component in network functionality (*7, 21, 33, 35, 51, 52, 60*). Moreover, recent work have shown that synaptic depression can desegregate different cusp bifurcations by weakening the strong synaptic effects between mutual excitatory loops (*46, 64*). Aside from that, by using plastic synapses, one can construct networks that are both balanced and near bifurcations (*14*). That is, mutual inhibition is not the only way to increase network capacity and flexibility; mutual inhibition is just one of many tools the brain could use to increase capacity.

A final direction for future work is to examine the best way to control networks with mutual inhibition. Several studies have suggested that networks near bifurcations are computationally useful (*14, 21, 61, 62*). Some studies use balanced inhibition to tune the network to be useful (*14*). Here, we wonder if this process can be used to control networks with mutual inhibition. If mutual inhibition is removed, are these networks still computationally useful?

To conclude, we have shown that mutual inhibition is key in expanding the functionality of the network. This is possible because mutual inhibition expands the number of cusp bifurcations in a network. Ultimately this helps elucidate why neural circuits in the brain are so flexible in their operation and are capable of such rich dynamics.

## Materials and Methods

### Spiking neural model and synaptic model

All simulations in this study are ran on flysim (*38*). Each individual neuron was modeled as a leaky integrate-and-fire neuron:

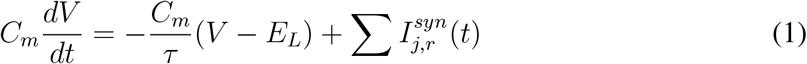

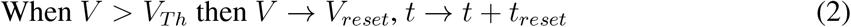

where *C_m_* is the capacitance, *V* the membrane potential, *τ* is the time constant, *E_L_* the reversal potential, and 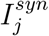 the current from the *j^th^* presynaptic neuron, and *r* represents the receptor. When the membrane potential reaches *V_Th_*, the neuron fires, and is promptly reset to *V_reset_* after an amount of time *t_reset_*. The parameters can be found in Table 1.

**Table 1:**
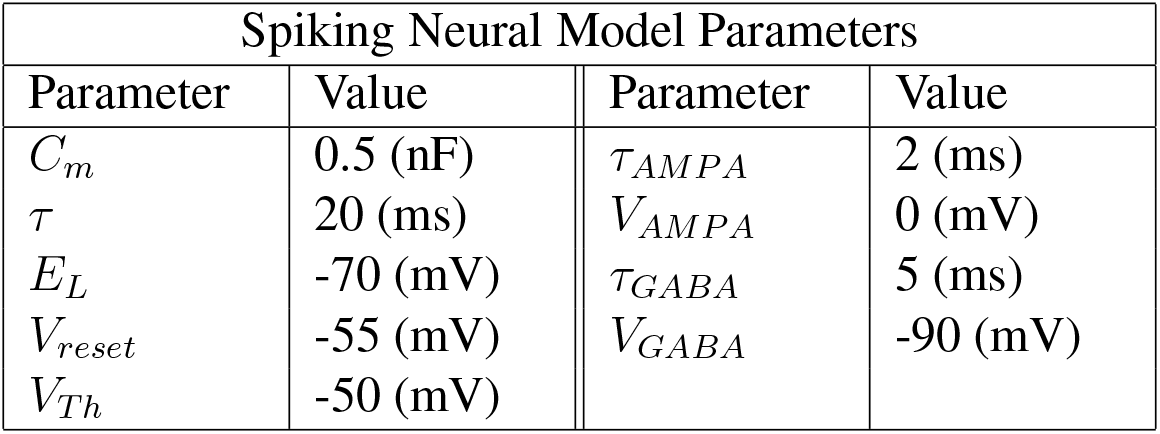
Parameters for neuronal model and synaptic model.

| Spiking Neural Model Parameters |  |  |  |
| --- | --- | --- | --- |
| Parameter | Value | Parameter | Value |
| $C_m$ | 0.5 (nF) | $\tau_{AMPA}$ | 2 (ms) |
| $\tau$ | 20 (ms) | $V_{AMPA}$ | 0 (mV) |
| $E_L$ | -70 (mV) | $\tau_{GABA}$ | 5 (ms) |
| $V_{reset}$ | -55 (mV) | $V_{GABA}$ | -90 (mV) |
| $V_{Th}$ | -50 (mV) | | |

Excitation is modeled as an exponentially decaying AMPA receptor, while inhibition is modeled as and exponentially decaying GABA receptor. More precisely, we have have excitatory and inhibitory current given by

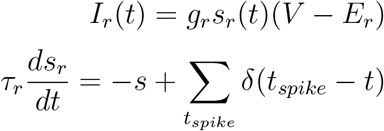

Here, *r* represents the receptor (*r* ∈ {*AMPA, GABA*}), *I* is the current of the receptor, *V* is the membrane potential, *E* is the reverse potential, *g* is the conductance of the receptor, *s* is the gating variable for the receptor ion channel. *s* is governed by a linear ODE with time constant *τ*, and is forced by a sum of dirac delta impulses ∑_*t_spike_*_ *δ*(*t_spike_* − *t*), where *t_spike_* is the timing of presynaptic spikes. It is important to note that *g_ee_* = *g_AMPA_* when both pre and post synaptic neurons are excitatory and *g_ei_* = *g_AMPA_* when presynaptic neuron is excitatiory and postsynaptic neuron is inhibitory. Likewise, *g_ii_* = *g_GABA_* when both pre and post synaptic neurons are inhibitory and *g_ie_* = *g_GABA_* when presynaptic neuron is inhibitory and postsynaptic neuron is exicatitory.

### Equilibrium points in microcircuits

It is conceivable that the functional repertoire of a motif is correlated to the abundance of its dynamical states. Here, we employ a statistical approach that provides us an estimate of the number of equilibrium points in relevant parameter space. We compared motifs that are specifically chosen to highlight the presence or absence recurrent loops, along with some randomly chosen feedforward motifs (Fig. 1b). After sweeping through the parameter space, we then estimated the number of equilibrium points in each trial by counting the number of clusters (S1a).

#### Parameter Space

For a four neuron motif, there are 12 possible synaptic connections. To lower dimensionality in parameters, we constrained synaptic weights with the same type of pre-synaptic and post-synaptic neuron to have the same value, except for threshold based filtering (Fig. 1a). Thus we have four types of synaptic weights: excitatory-to-excitatory (*g_ee_*), excitatory-to-inhibitory (*g_ei_*), inhibitory-to-inhibitory (*g_ii_*) and inhibitory-to-excitatory (*g_ie_*). For each motif, we ran simulations over 18900 trials, where each trial has a different synaptic weight parameter set *g_ee_, g_ie_, g_ii_* (Table S1).

Aside from synaptic weights, we also iterated through different underlying bias currents in the neurons. The bias currents adjust the threshold for firing, allowing inhibitory neurons to fire in the absent of excitatory input (Table S1). This ensures that we capture the dynamic states produced by the inhibitory neurons. For each trial, a brief injected current goes into different pairs of neurons, for example one trial would stimulate *E*_1_ only, and another would stimulate *E*_1_ and *I*_1_ simultaneously. This is ran through all possible pairs. The amplitude of the stimulus changes as well. Both stimulating different pairs as well as varying the stimulus amplitude ensures a wider exploration of the parameter space, thereby allowing us to find more equilibrium points. Each brief pulse of positive stimulation is separated by a negative reset signal. These are shown in S1b. Note that this method may not capture every single equilibrium point present, nor will it be able to detect other stable structures such as limit cycles. Nevertheless, the parameter sweeping still provides valuable information regarding the complexity of the dynamical states of the system.

#### Clustering

To analyze the amount of equilibrium points present in each trial, we look at the firing rate trace of the motif across the four-dimension space (*e*_1_, *e*_2_, *i*_1_, *i*_2_). We used the elbow method to obtain the amount of clusters in the firing rate trace of the trial (*65*). The elbow method detects the number of clusters by iterating through possible number of clusters *k* and performing K-means clustering, and then evaluating each guess *k* with its standard deviation, which is 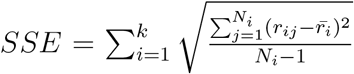. By plotting *SSE* as a function of *k*, we see that sometimes the plot has a sharp drop in *SSE* followed by a shallow decrease when *k* becomes large. The sharp drop is cleverly termed “the elbow” (Fig. S1a). Such a sharp drop usually indicates that the number of clusters assigned has hit the correct amount. If no elbows are present, then the data set most likely only has one cluster. A criteria is needed to determine “how sharp” a drop can be considered an elbow, and this criteria is set so that the slope of the “arm” must be seven times the slope of the “forearm”.

We purposely neglected the transient activities 100ms after a positive stimulation pulse or a negative reset signal in order to neglect “fake” clusters (i.e. clusters that are present, but are not true equilibrium points). Finally, the total number of equilibrium points in all possible parameter sets, which total up to 18900 trials, are calculated for each motif for comparison.

### CPG in microcircuits

Another important computation for small motifs is the ability to generate oscillations under constant input. Similar to how the equilibrium points were counted, for CPGs, the parameter space was scanned across suitable values of *g_ie_* and *g_ei_*. For each trial, we then determine whether the motif under said parameter set was capable of oscillation. The ISI for each trial was calculated, and rounded to the nearest 0.001 (s). Since there is no noise in the system, the periodic firing of a neuron being driven by a constant current has a near perfect period (where the slight differences were accounted for by the rounding procedure), and thus the ISI would consist of one single value. The choice of rounding, 0.001 (s), is hence chosen so that when receiving a constant bias current with no noise, the ISI will be 1 single value, as it should be. For any rounding numbers smaller than that, for example 0.0005 (s), such trials will yield 2 or more ISI values, which is due to time step choices of the simulator and does not really reflect whether the motif is in a CPG mode. For CPGs, the ISI would contain two or more values. The counts for trials with CPGs were calculated for each motif.

### Decision Making and Bistability

To further illustrate how feedback and mutual inhibition differs in terms of functions, we focused on two functions in particular: decision making and bistability. We swept through the relevant parameter space, and observed whether motifs with or without mutual inhibition could perform these two functions. Once again, to lower the dimensionality of the parameter space, we constrained synaptic weights with the same type of pre-synaptic and post-synaptic neuron to have the same value. For bistability, since it happens within the excitatory subsystem, the most crucial synaptic weight is *g_ee_*. For decision making in feedback inhibitory networks, the important synaptic weight is *g_ie_*, since strong *g_ie_* allows the circuit to turn off the unfavorable excitatory neuron. For decision making in mutual inhibitory networks, the key synaptic weight is *g_ii_*. Therefore, for the purpose of this analysis, we will hold *g_ei_* constant and sweep through relevant parameter regimes of *g_ee_, g_ie_* and *g_ii_*. The parameters are listed in Table S2.

For each point (*g_ee_*, *g_ie_*, *g_ii_*) in the parameter space, two trials with different input protocols are given to the network. The bistable protocol gives a 100 ms positive pulse to one of the excitatory neurons, resets the circuit using a strong negative pulse, and repeats this process for all of the excitatory neurons present within the network (Fig.S2). If the network is capable of bistability, then the pulse to one of the excitatory neurons should be able to turn the network to the ON state and remain there until the resetting pulse comes in. Therefore, if the circuit stays on after *E*_1_ is excited, and also stays on after *E*_2_ is excited, then it is classified as being able to perform the switch function.

The decision protocol iterates through all possible pairs of excitatory neurons, where for each time period each pair is given an asymmetrical input (Fig. S2). If the network decides, then there will be a difference in firing rate between a pair of neurons at that time period. The condition for decide is that the “winning” neuron (if one neuron’s time-averaged firing rate is 10 Hz larger than the other, it is considered a winner) must switch as the relative input strength switches. For instance, if *E*_1_ wins when input 1 is larger than input 2, then *E*_2_ must win when input 2 is larger than input 1. For motif B, decision is made between the excitatory neurons, hence the aforementioned analysis is performed on the two excitatory neurons. The decision is made between the inhibitory neurons for motif A, hence the decision analysis here is performed on *I*_1_ and *I*_2_. Each point (*g_ee_, g_ie_, g_ii_*) will be capable of either bistability, quasi-decision, decision, both, or none. The results are then drawn into a heat map, where broader regions of “both functions present” means that it is more possible for the two functions to co-exist.

### Dynamical analysis

For further understanding of the statistical results, we reduced the CRIREL system into a two dimensional model given by the equations

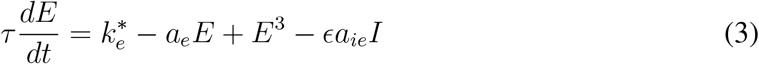

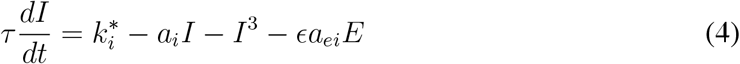

where, 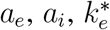, and 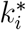 are the parameters of the cusp bifurcations. *ϵa_ie_* and *ϵa_ie_* are the coupling strengths between each subsystem. The full derivation of the reduced model is given in the Supplementary Note. In this paper we only consider small *ϵ*, and consider large *ϵ* out of the scope of this paper. The reduced model allows us to conceptually understand what is occurring dynamically in the spiking neural network. Furthermore, the intuition gained from the reduced model is invaluable, as any function found in the reduced model can also be found in the spiking model for an appropriate set of parameters. Note that this does *not* work in the converse direction, meaning there could be functions in the spiking model that are not in the reduced model. This is especially apparent when *ϵ* is not small.

As a broad overview, we first begin by showing that the mutual excitatory loop and the mutual inhibitory loop each have a cusp bifurcation. To contrast this, we also showed that feedback inhibition cannot undergo a cusp bifurcation. Next, we coupled the two circuits together, and show that both cusps are maintained. Finally, we worked out the reduced model’s simple cubic form.

### CRIREL function simulations

The reduced model allowed us to predict the different functions the CRIREL circuit may be capable of performing. These functions are actualized by simulation. While the reduced model is general for the whole class of CRIREL circuits, the results we have shown are simulated on one particular circuit (motif A in Fig. 1b), with the exception of threshold-based filtering, which is simulated on motif H (Fig. 6a). The parameters for simulation are listed in Table S3, and the stimulation are shown in the results section. Parameter searches are illustrated in detail below.

#### Switch, Toggle and Anti-Toggle

The robustness of the functions determines how realistically it can be implemented in noisy biological systems. For switch, toggle and anti-toggle, we swept through appropriate regimes to showcase its working parameter range (Fig. 3c, f, Fig. 4f). A pulse of a specific amplitude and duration was given to the circuit, in the same manner as displayed in Fig. 3a, d and Fig. 4f respectively.

If the circuit is turned on – defined as mean firing rate larger than zero in the designated time period (200-400 ms) – and off – defined as mean firing rate equal to zero after 500 ms, then the circuit is deemed capable of performing switch/toggle within said parameter set. Toggling is furthermore sensitive to the precise timing of the second pulse near the transition to toggling functionality, which means that within certain parameter sets, the behavior of toggling is not robust when the onset or offset pulse timing varies. Therefore, the onset of the pulse is varied from 500-505 ms, and the circuit is only labeled as capable of performing toggling if it toggles for all 6 onset times.

Similarly, a parameter set is defined as performing anti-toggle if the designated neuron fires after the duration of the input ends, and the other gradually returns to silence (evaluated by seeing if the designated neuron has a higher firing rate than the other).

#### sCPG and aCPG

For central pattern generators, the ability to adjust its firing period is essential (Fig. 3i, Fig. 4i). This can be achieved by modifying its input into the inhibitory neurons. Again, the protocol mimics those of Fig. 3g and Fig. 4g respectively. The ISI of the inhibitory neurons is calculated by averaging all the ISIs within one simulation trial, while disregarding the initial ISI.

#### Decision making

The psychometric function of the decision making circuit is shown to quantify its accuracy and reaction time (Fig. 4c). The input difference 2*c* (%) is defined similar to those of (*6*), where the input to one population (in our case, one inhibitory neuron) is 1 + *c*, and the input for the other neuron is 1 − *c*, and *c* ranges from 0 to 1. Essentially, the larger the input difference, the easier the decision is to make. The input lasts for 100 ms. The accuracy is defined as the number of correct trials over total trials, and in cases where no decision is reached, the winning population is randomly assigned. The reaction time is defined as the time from which the stimulus begins to the time where one population reaches the threshold firing rate of 200 Hz. In cases with no decisions, the reaction time is set at 2 s. The mean reaction time is then calculated by averaging over all 500 trials.

#### Threshold-based filtering

By tuning the bias current of *I*_1_, the threshold of the filter can be adjusted. The stimulation used here is identical to those of Fig. 6b. The input “passes” the threshold when the maximum firing rate of *E*_2_ reaches 100 Hz within 100 ms after stimulation. Note that the maximum passed input amplitude appears to be discrete because the input used in Fig. 6b is discrete itself.

#### Coincidence and Anti-coincidence detectors

Similar to how we tested the working parameter range for switch and toggle, we swept through different pulse amplitudes and timing differences for the coincidence and anti-coincidence detectors. A working parameter set is defined as the circuit firing according to its truth table (1 for firing rate larger than zero, and 0 for firing rate equal to zero).

### Large random network

To determine whether the flexibility of mutual inhibition is preserved in larger networks, we built a larger network and investigated a specific function, memory. To construct the large network, we used 100 LIF neurons with the same membrane parameters as the microcircuits. The excitatory population had 75 neurons, while the inhibitory population had 25 neurons. The parameters for connection probability and synaptic weights are given in Table S4.

Each excitatory neuron was given a random 500 ms pulse with an amplitude drawn from an normal distribution with mean *μ* = 3 and variance *σ*^2^ = 3 in addition to an underlying bias current of −5 nA. After the pulse ends, we waited for an extra 1 s before giving a reset signal of −20 nA to the excitatory neurons. Through this protocol, if the network is capable of sustaining memory, then it will settle into a random equilibrium point after the positive pulse is removed, and before the reset signal arrives. Three different types of entropy are calculated to analyze the variety of memory states in the network.

#### Entropy

For a discrete random variable *X*, where there are *n* potential outcomes *x_i_*, …*x_n_* with associated probability *P*(*x*_1_), …*P*(*x_n_*), its entropy is

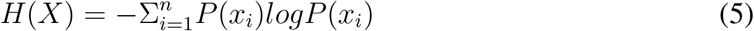

#### Entropy of words, *H_word_*

After the pulse is turned off and before the reset signal arrives, each neuron *i* will have a different time-averaged firing rate 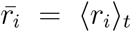, which represents the “state” the neuron is in for that particular equilibrium point. Together, these firing rates can form a sequence, i.e. *r*_1_, *r*_2_, *r*_3_, …*r*_1_00, which we call a “word”. Each trial will produce one such word. If the memory network has multiple equilibrium points, then the variation of words would be large, and the entropy resulting from such a distribution of words, *H_word_*, would be large.

When calculating the entropy, what we actually used is the quotient of 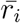 over 20. This is similar to grouping the firing rate into different bins, where the firing rates within the same bin is assumed to carry the same amount of information. The choice of having a 20 Hz bin is arbitrary – qualitatively similar results can be obtained by other bins, such as 30 Hz and 40 Hz as well.

#### Entropy of symbols, *H_symbol_*

A word *r*_1_, *r*_2_, *r*_3_, …*r*_100_ itself is a distribution of firing rates. The entropy of this distribution is called *H_symbol_*, which reflects how widely distributed these firing rates are. For the word produced by each trial, its entropy of symbols is calculated, and then averaged across trials.

#### Entropy of binary symbols, *H_binary_*

Instead of representing each neuron by its firing rate, we instead observe whether the neuron fires or not. If it fires, the neuron is given a label *l_i_* = 1, and *l_i_* = 0 otherwise. This again forms a distribution across the neuron population, *l*_1_, *l*_2_, *l*_3_, …*l*_100_, and its associated entropy is called *H_binary_*.

## Supporting information

augmenting_flexibility_supplementary

## Acknowledgments

We thank Gabrielle Gutierrez and for discussion and feedback on the figures, Youngmin Park for useful discussions on the mathematical proofs presented in the study, and Fred Rieke and Peter J. Thomas for feedback on the manuscript. This work is partially supported by the Featured Areas Research Center Program within the framework of the Higher Education Sprout Project, a joint fund from the Ministry of Education (MOE) and the Ministry of Science and Technology (MOST) in Taiwan. This work was also partially supported by Ministry of Science and Technology grant number 109-2218-E-007-021-in Taiwan.

## Supplementary materials

Materials and Methods

Supplementary Text

Figs. S1 to S3

Tables S1 to S4

References (*4–10*)

### Dynamical Analysis

A reduced model was used to gain a deeper understanding of the dynamics of the CRIREL circuit. Here we derive the reduced model that we use to organize our results. The reduced model allows us to conceptually understand what is occurring dynamically in the spiking neural network. Furthermore, the intuition gained from the reduced model is invaluable, as any function found in the reduced model can also be found in the spiking model for an appropriate set of parameters. This is a consequence of central manifold reduction. Note that this does not work in the converse direction, meaning there could be functions in the spiking model that are not in the reduced model.

As a broad overview, we first begin by showing that the mutual excitatory loop and the mutual inhibitory loop each have a cusp bifurcation. Then we prove that feedback inhibition cannot undergo a cusp bifurcation. Next, we show that when coupled together, both cusps are maintained. Then finally, we work out the reduced model’s simple cubic form.

### Decoupled mutual excitation and mutual inhibition

We begin with a simple firing-rate model for the mutual excitation loop:

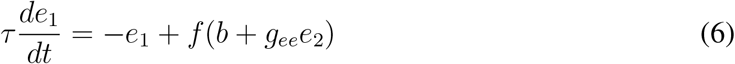

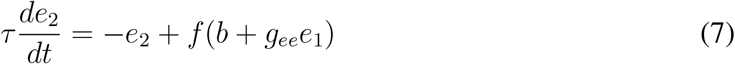

and mutual inhibition:

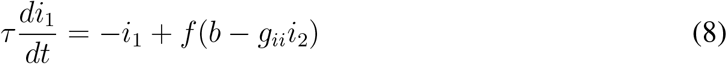

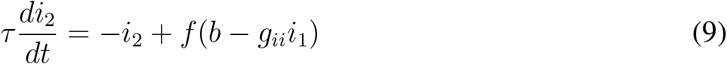

Here, *e*_1_, *e*_2_, *i*_1_, and *i*_2_ are the firing rates of the neurons, and *τ* is a time constant. Our two bifurcation parameters are the bias current *b* and the synaptic weight *g_ee_* or *g_ii_*, depending on the type of mutual connection. Because the LIF spiking model we use is a class 1 excitable neuron (*54*) (it has a continuous input frequency curve), we assume that *f*(*x*) is monotonic, i.e. *f′*(*x*) ≥ 0.

To begin, we need to find the cusp bifurcation point. A necessary condition of the bifurcation point is the point at which one of the eigenvalues of the Jacobian is 0. Thus we calculate the Jacobians for our system at the equilibrium point (denoted with a * super-script) and find:

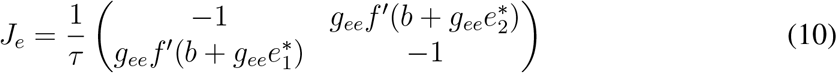

and for inhibition:

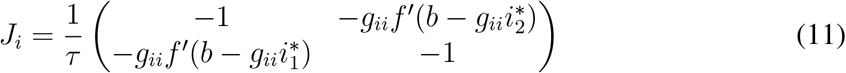

Note here that *f′* is positive. Simplifying the notation and letting 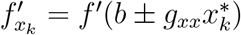

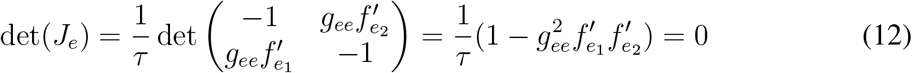

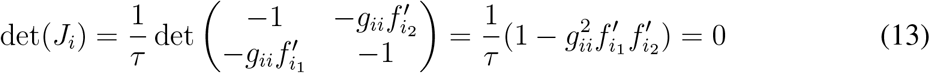

Thus there will be at least a saddle node bifurcation whenever 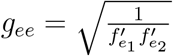 or 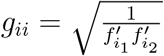.

Note that for feedback inhibition we cannot even have a saddle-node bifurcation. If we examine the firing-rate model

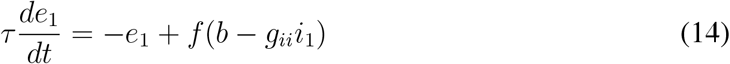

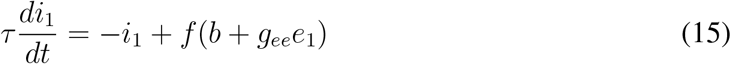

we see that the Jacobi an can never have a 0 eigenvalue.

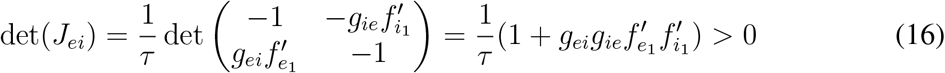

This is because all terms are restricted to be positive. Thus feedback inhibition can only have one stable fixed point.

The next step here to prove that we have a cusp bifurcation, and not just a saddle-node bifurcation, is to prove that the quadratic part of the system degenerates. To do this we need to find a center manifold, expand along that manifold, and show that the quadratic term can be zero. We begin by finding the center manifold for the two systems. This is a one dimensional surface such that *e*_2_ = *M_e_*(*e*_1_) and *i*_2_ = *M_i_*(*i*_1_).

We can calculate the manifold by noting that

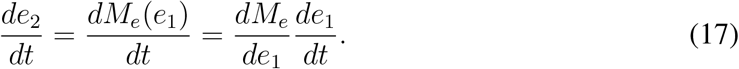

and

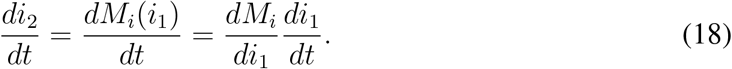

Solving for *M_i_* and *M_e_* gives us

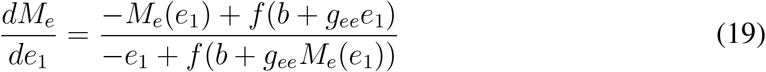

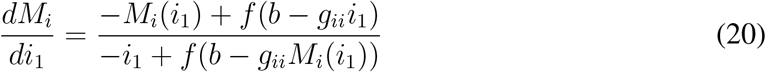

In this particular case the excitatory manifold is easier to algebraically determine the excitatory manifold, so we will proceed only with the math for the excitatory subsystem here. The inhibitory center manifold *M_i_* must be solved for numerically.

The excitatory manifold is solvable with the ansatz *e*_2_ = *e*_1_ = *M_e_*(*e*_1_). Plugging this in gives us,

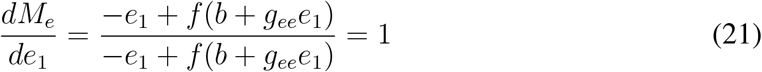

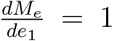 can be solved very easily by separation of variables, giving us *M_e_*(*e*_1_) = *e*_1_, thereby proving that the ansatz is valid.

Continuing onward, we have the dynamics constrained on a one dimensional manifold *e* ≔ *M_e_* and *M*(*i*) ≔ *M_i_*. Here, we have switched to *e* and *M*(*i*) to emphasize the fact that *i* is no longer in the same coordinates as *i*_1_ and *i*_2_.

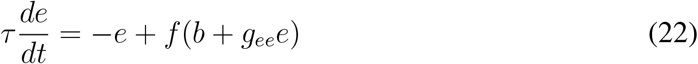

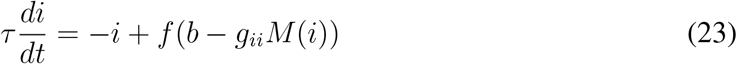

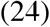

We can now expand this using a Taylor series.

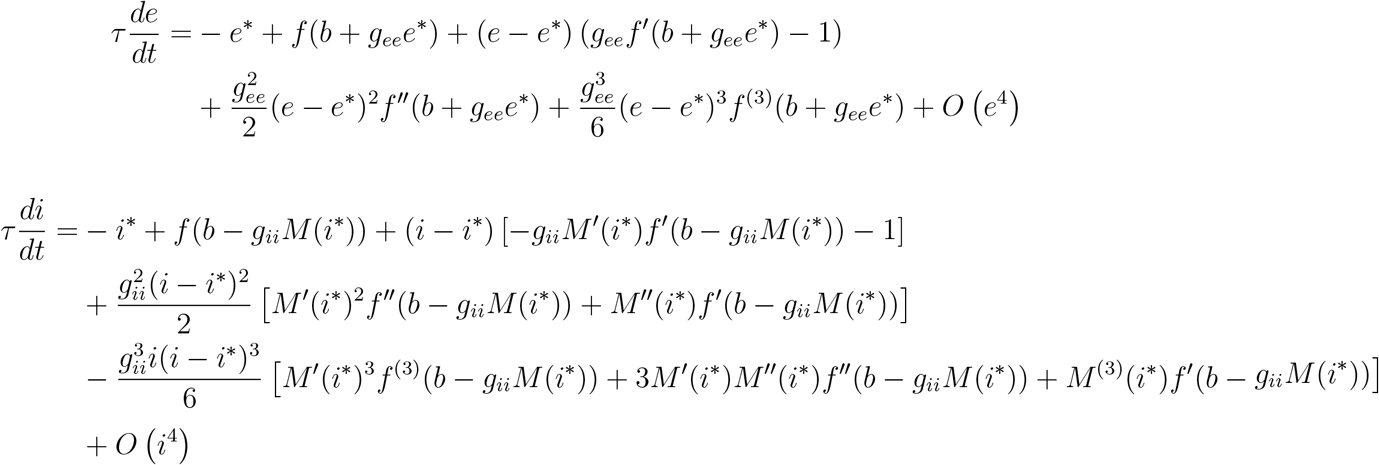

Using a change of variables, we can convert the above equations into two depressed cubics giving us

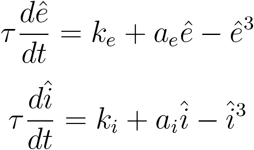

where

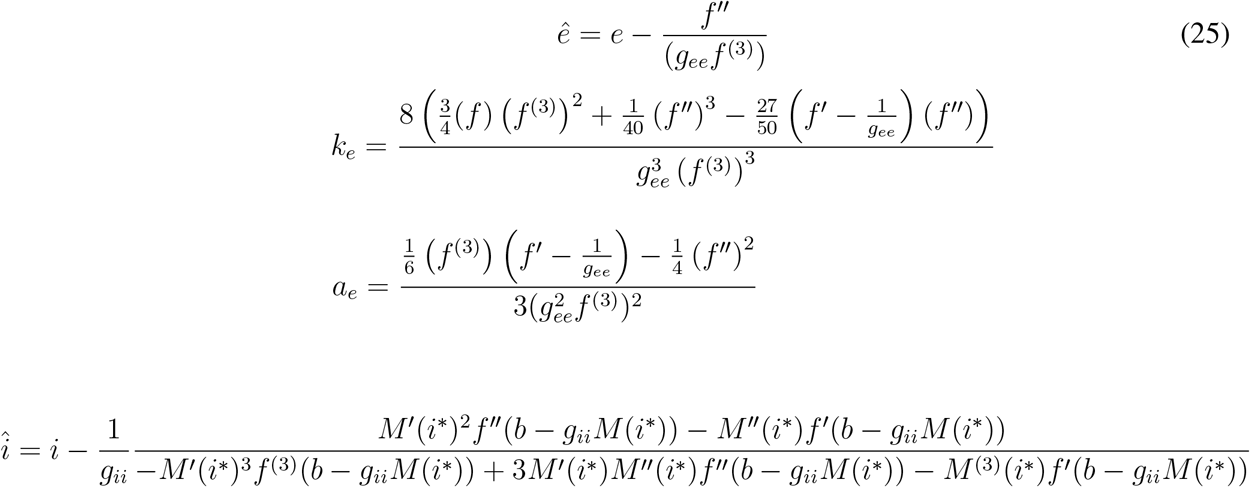

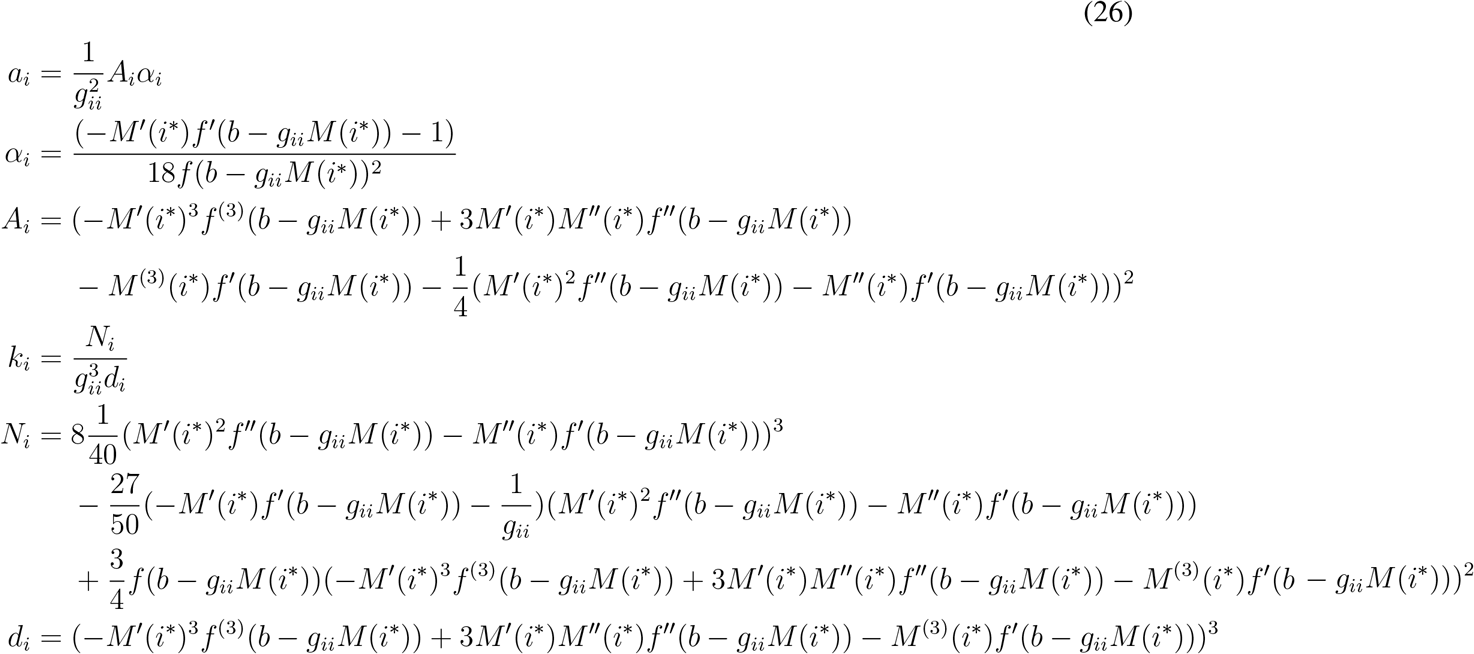

Whenever *k_e_*, and *a_e_*, or *k_i_*, and *a_i_* are 0, then the two systems undergo a cusp bifurcation. While the exact values of the cusp bifurcation is complicated, especially in the case of the inhibitory system, it still can undergo the bifurcation.

### Coupling mutual excitation with inhibition

Now that we have calculated the normal form for both of subsystems, we can couple them together. We don’t need to explicitly use the normal form at first. Rather its mere existence will be used later. We begin by introducing the coupling terms *ϵ*. The corresponding equation becomes

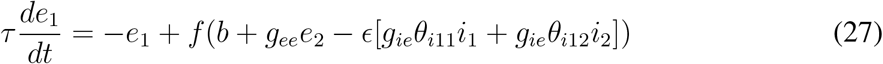

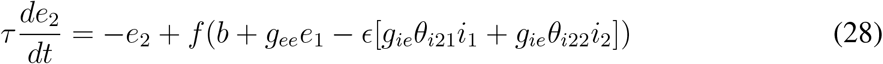

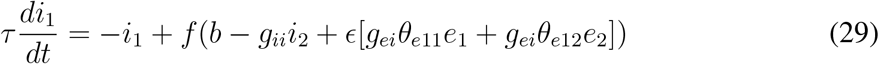

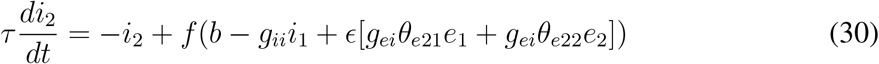

Here, 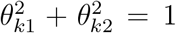 and *θ_kx_* > 0. This allows the coupling between the connections to be arbitrary. We can think of *θ* as a way to parameterize the asymmetries in synaptic weights between the coupling.

Next, by expanding the function *f* as a Taylor series with respect to *ϵ* centered at *ϵ* = 0, and then disregarding order *ϵ*^2^ and above, we get

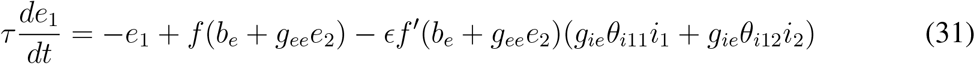

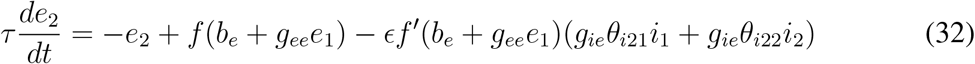

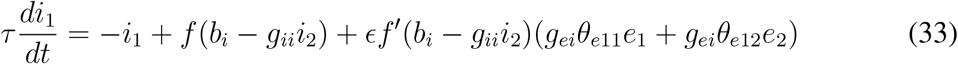

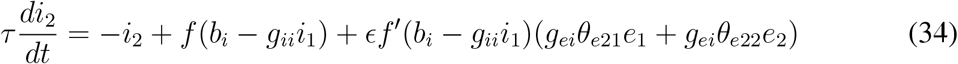

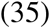

We can then make the substitution for the stable attractive manifolds we made above such that *e* = *e*_2_ = *M*(*e*_1_) = *e*_1_ and *i* = *i*_2_ = *M*_i_(*i*_1_).

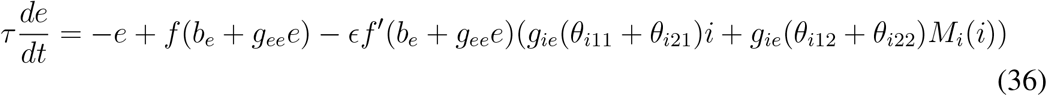

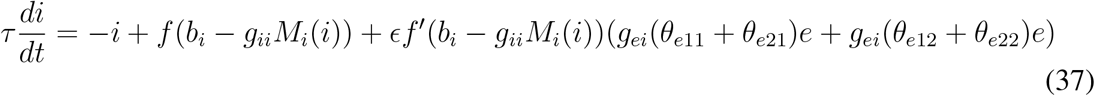

For ease of notation let *θ*_*i*1_ = *θ*_*i*11_ + *θ*_*i*21_ and *θ*_*i*2_ = *θ*_*i*12_ + *θ*_*i*22_, and *θ*_*e*1_ = *θ*_*e*11_ + *θ*_*e*21_ and *θ*_*e*2_ = *θ*_*e*12_ + *θ*_*e*22_ We can now expand the equation in terms of *e* and *i* giving us

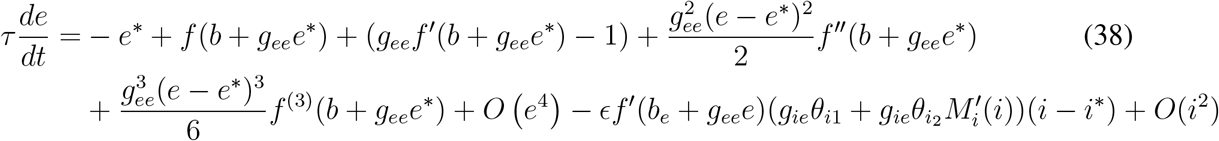

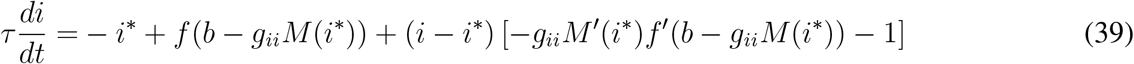

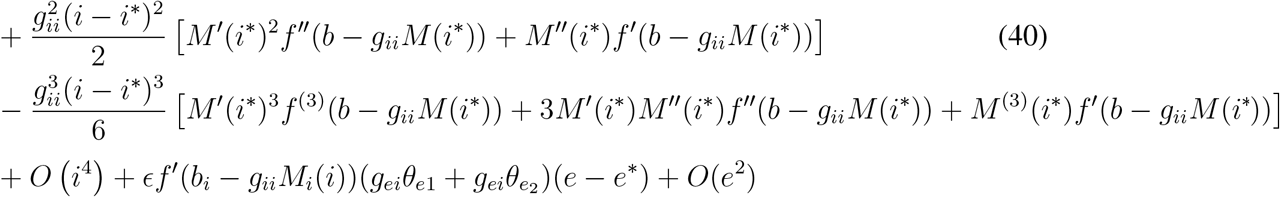

Recall we can create a depressed cubic in the uncoupled system using equations 25 and 26 to get

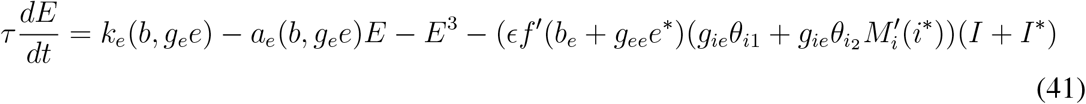

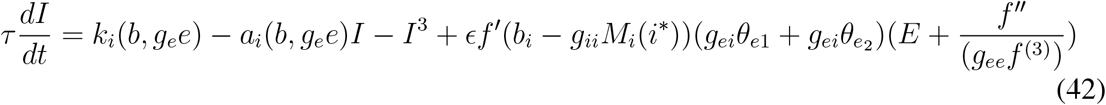

where 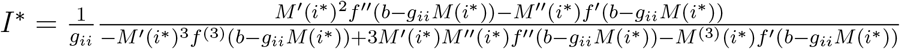 from equation 26. To clean everything up a bit, we can recast our equations as

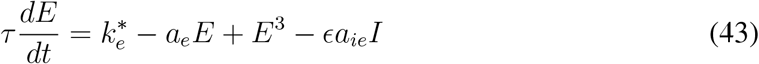

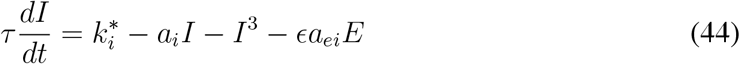

where *a_ie_* = *g_ie_θ*_*i*1_ + *g_ie_θ*_*i*2_*M′*, *a_ei_* = *g_ei_θ*_*e*1_ + *g_ei_θ*_*e*2_, 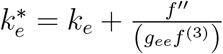 and 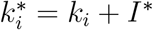.

### Supplementary Figures

**Figure S1:**
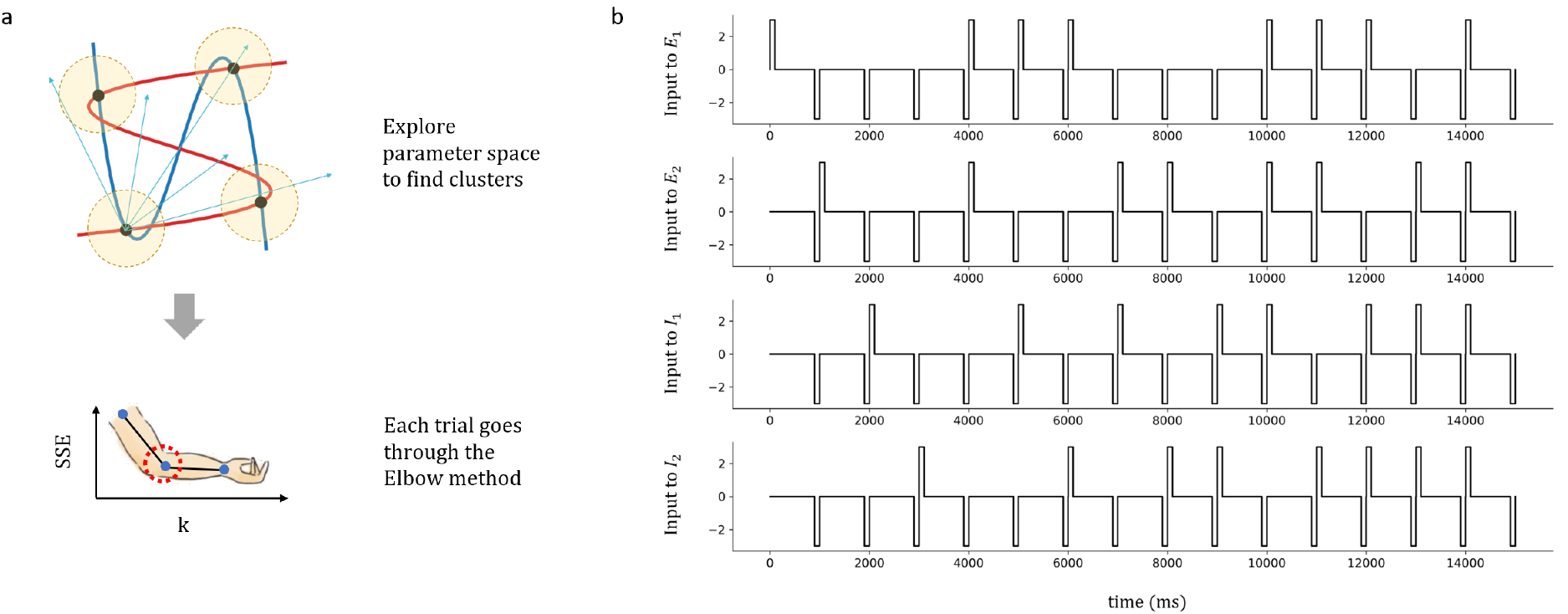
Protocol for statistical analysis of equilibrium points. (a) *Top:* Schematic diagram of attractors in a motif. Each attractor has its own basin of attraction, so that once the system is pushed near the attractor, it stays near the attractor. The protocol is designed to explore different state spaces so as to find as many attractors as possible. *Bottom:* The method we used to determine the number of equilibrium points is the elbow method. Specifically, when the algorithm hits the number of correct clusters *k* in the data, it will result in an elbow-like dip in *SSE*, as shown in the graph. (b) The current stimulation given to the four neurons in one trial. The duration for the positive pulse and reset pulses are both 100 ms, while the interval in between the pulses are 800-900 (ms) – the precise interval value doesn’t matter, as long as they are longer than the transient activities.

**Figure S2:**
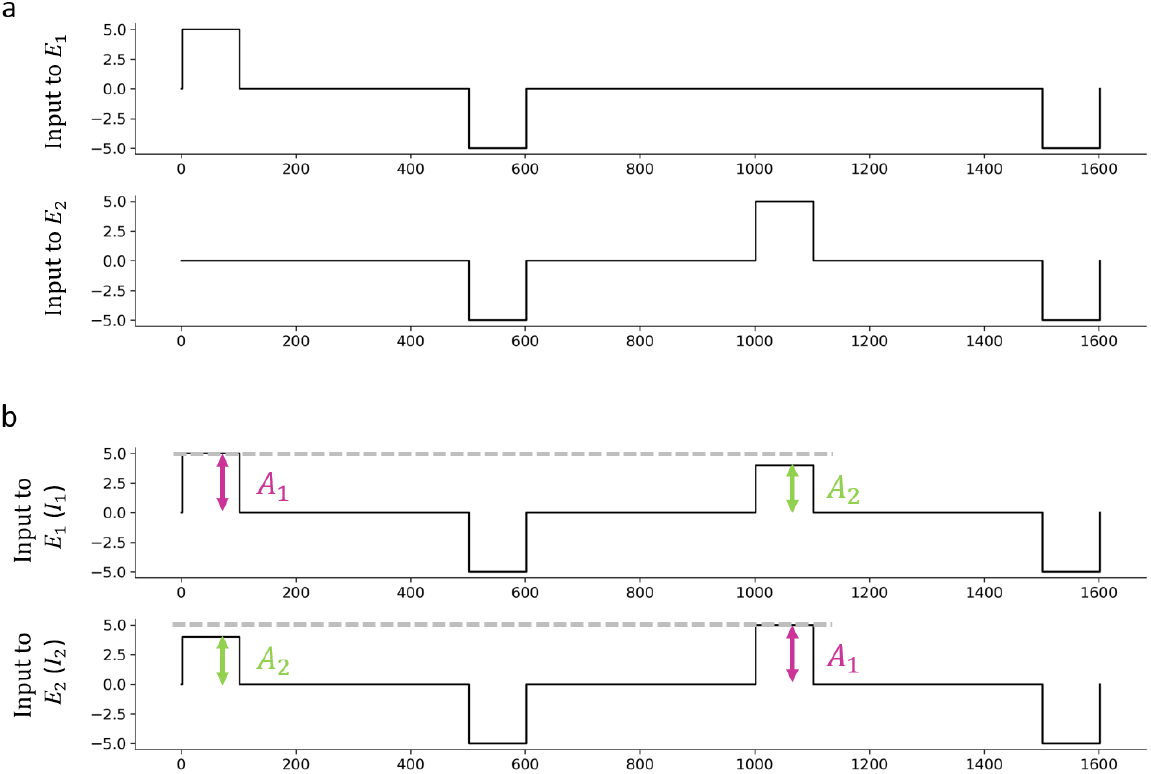
Input protocol for decision making and switch. (a) The input protocol for switch. The duration of the pulses are 100 ms, and the interval in between is 400 ms. The strength of the positive and reset pulses is 5 and −5 nA, respectively. (b) The input protocol for decision making. Note that *A*_1_ and *A*_2_ have different amplitudes (*A*_1_ = 5 nA and *A*_2_ = 4 nA), representing how the input signal strength is different for the two neurons. The reset pulse is −5 nA. No noise is included in this simulation, but including noise does not change the results qualitatively.

**Figure S3:**
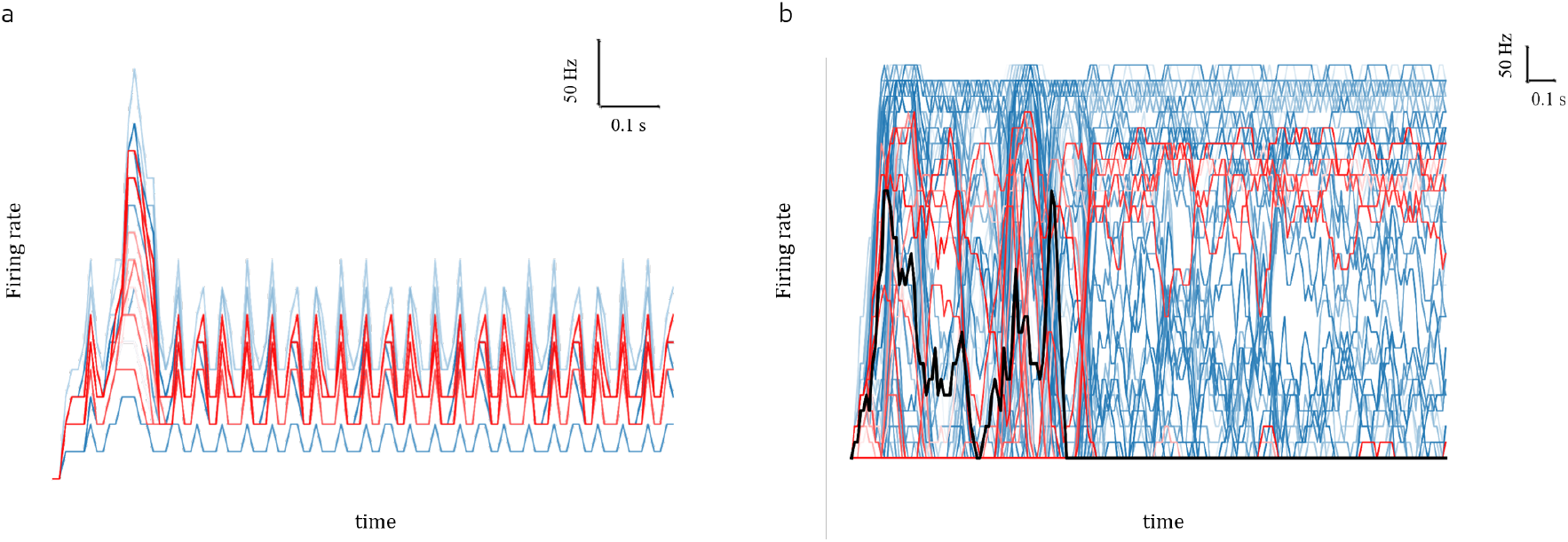
CPG and toggle in large random network. (a) Firing rate of neurons in the network performing CPG, blue is excitatory and red is inhibitory. (b) Firing rate of neurons in the network performing toggle, blue is excitatory and red is inhibitory. Black is one particular neuron used to showcase toggle more clearly. The parameters for toggle is: *g_ee_* = 25 (*μS*), *g_ei_* = 15 (*μS*), *g_ie_* = 150 (*μS*), *g_ii_* = 50 (*μS*), *b_E_* = 2.5 (*nA*), *b_I_* = 0 (*nA*). The parameters for CPG is: *g_ee_* = 25 (*μS*), *g_ei_* = 25 (*μS*), *g_ie_* = 50 (*μS*), *g_ii_* = 25 (*μS*), *b_E_* = 1 (*nA*), *b_I_* = 0 (*nA*).

### Supplementary Tables

**Table S1:**
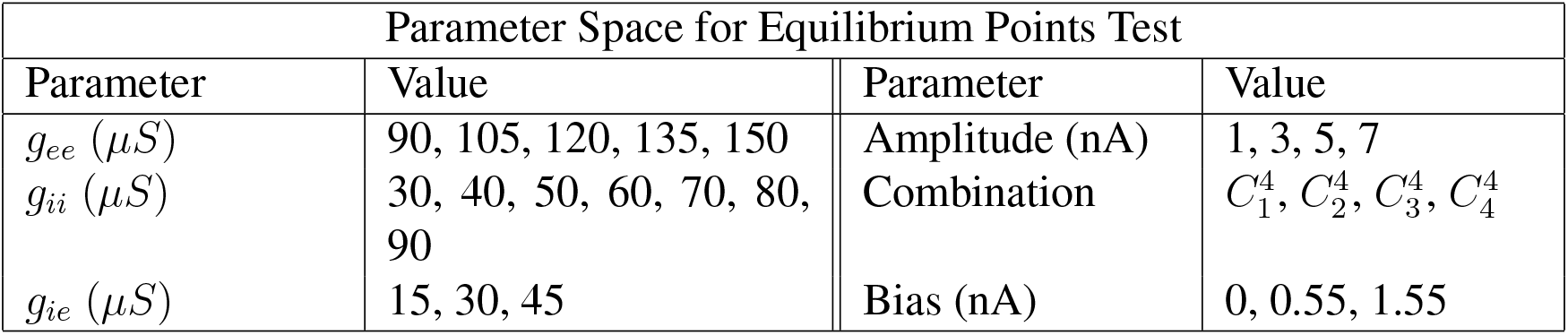
Parameter space for determining the equilibrium points of the motifs. *g_ei_* is fixed at 30 *μS*. Amplitude represents the amplitude of the positive pulses given in units of nA. The amplitude for the reset pulses are fixed at −3 nA. Combination reflects the different combinations of neurons that are given the pulse. For example, at a given time period, neurons *E*_1_ and *E*_2_ might receive current pulses while neurons *I*_1_ and *I*_2_ do not. This is iterated through all different combinations. Bias represents the underlying bias currents that all neurons receive.

**Table S2:**
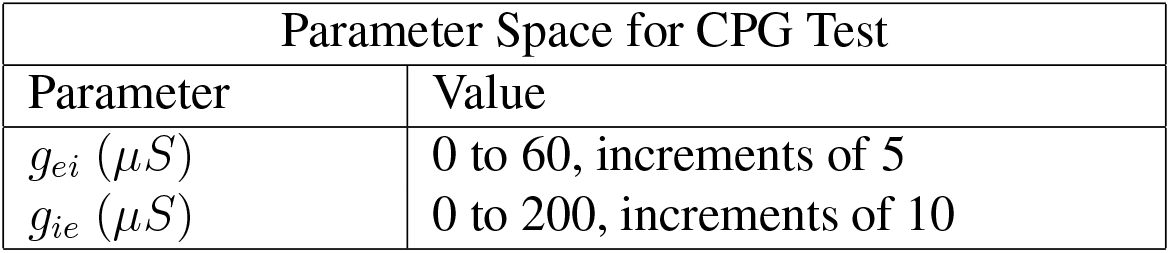
Parameter space for determining the CPG of the motifs. *g_ee_* is fixed at 65, *g_ii_* at −5, the bias current for excitatory neurons is *b_E_* = 0.5, and the bias current for inhibitory neurons is *b_I_* = −0.5.

**Table S3:**
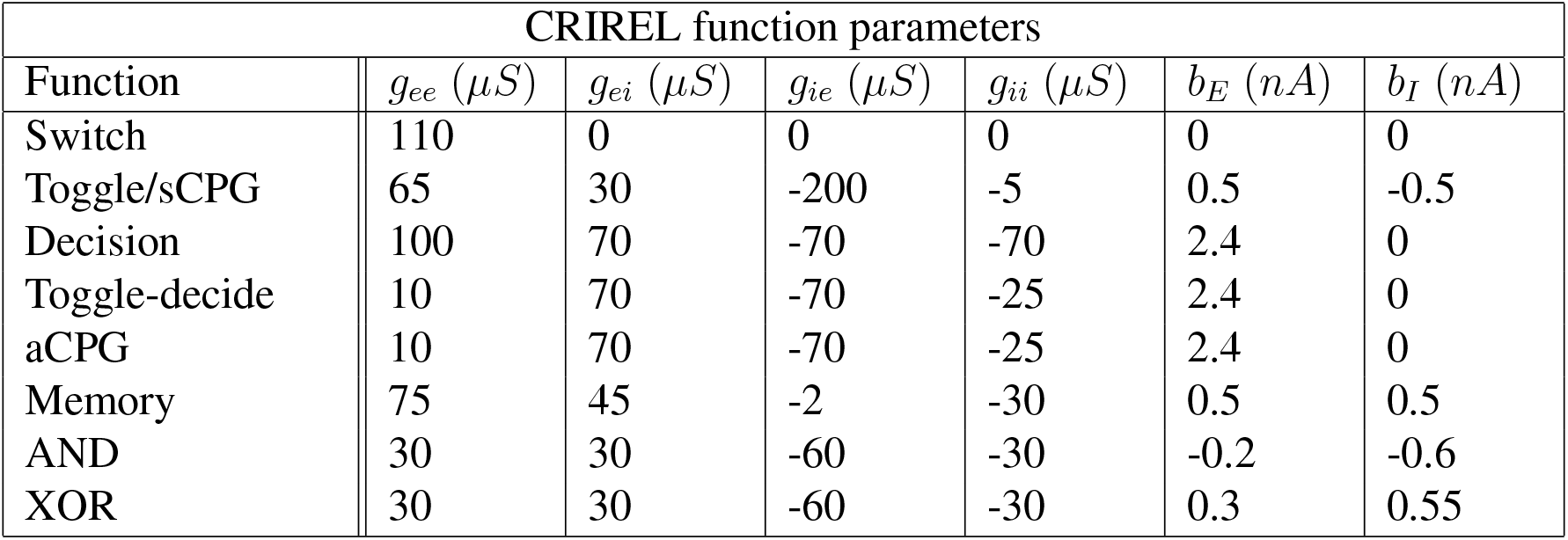
The parameters used to produce the CRIREL circuit functions in the figures. *b_E_* is the bias current for excitatory neurons, and *b_I_* is the bias current for inhibitiory neurons. The parameters for the neuronal and synaptic model follows the parameters given in the methods section unless otherwise specified. The parameters for amplitude-based decision is more complicated, and is listed here instead: *g_ee_* = 50 (*μS*), *g_ie_*=-4000 (*μS*), *g*_*E*_1_*toI*_2__ = 25 (*μS*), *g*_*E*_1_*toI*_1__ = 20 (*μS*), *g*_*I*_1_*toI*_2__ = 70 (*μS*), *g*_*I*_2_*toI*_1__ = 30 (*μS*). Also, the membrane time constant *τ* is 80 (ms) and capacitance *C* is 1 *nF* for *I*_2_.

**Table S4:**
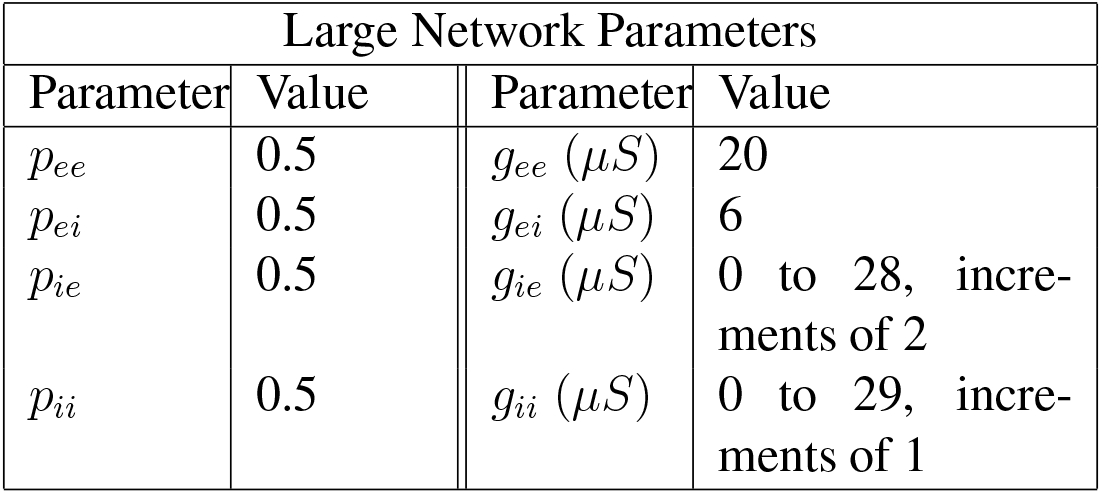
Connection probability and synaptic weight of large network. *p_ij_* represents the connection probability of a neuron of type *j* to type *i*, where *i, j* ∈ *E, I*. *g_ij_* represents the synaptic weight of a neuron of type *j* to type *i*.

