## Supplementary material for "Augmenting Flexibility: Mutual Inhibition Between Inhibitory Neurons Expands Functional Diversity": augmenting_flexibility_supplementary

### Dynamical Analysis

A reduced model was used to gain a deeper understanding of the dynamics of the CRIREL circuit. Here we derive the reduced model that we use to organize our results. The reduced model allows us to conceptually understand what is occurring dynamically in the spiking neural network. Furthermore, the intuition gained from the reduced model is invaluable, as any function found in the reduced model can also be found in the spiking model for an appropriate set of parameters. This is a consequence of central manifold reduction. Note that this does *not* work in the converse direction, meaning there could be functions in the spiking model that are not in the reduced model.

As a broad overview, we first begin by showing that the mutual excitatory loop and the mutual inhibitory loop each have a cusp bifurcation. Then we prove that feedback inhibition cannot undergo a cusp bifurcation. Next, we show that when coupled together, both cusps are maintained. Then finally, we work out the reduced model's simple cubic form.

### Decoupled mutual excitation and mutual inhibition

We begin with a simple firing-rate model for the mutual excitation loop:

$$\tau \frac{de_1}{dt} = -e_1 + f(b + g_{ee} e_2) \tag{6}$$
$$\tau \frac{de_2}{dt} = -e_2 + f(b + g_{ee} e_1) \tag{7}$$

and mutual inhibition:

$$\tau \frac{di_1}{dt} = -i_1 + f(b - g_{ii} i_2) \tag{8}$$
$$\tau \frac{di_2}{dt} = -i_2 + f(b - g_{ii} i_1) \tag{9}$$

Here, $e_1$, $e_2$, $i_1$, and $i_2$ are the firing rates of the neurons, and $\tau$ is a time constant. Our two bifurcation parameters are the bias current $b$ and the synaptic weight $g_{ee}$ or $g_{ii}$, depending on the type of mutual connection. Because the LIF spiking model we use is a class 1 excitable neuron (*54*) (it has a continuous input frequency curve), we assume that $f(x)$ is monotonic, i.e. $f'(x) \geq 0$.

To begin, we need to find the cusp bifurcation point. A necessary condition of the bifurcation point is the point at which one of the eigenvalues of the Jacobian is 0. Thus we calculate the Jacobians for our system at the equilibrium point (denoted with a $^*$ super-script) and find:

$$J_e = \frac{1}{\tau}\begin{pmatrix} -1 & g_{ee} f'(b + g_{ee} e_2^*) \\ g_{ee} f'(b + g_{ee} e_1^*) & -1 \end{pmatrix} \tag{10}$$

and for inhibition:

$$J_i = \frac{1}{\tau}\begin{pmatrix} -1 & -g_{ii} f'(b - g_{ii} i_2^*) \\ -g_{ii} f'(b - g_{ii} i_1^*) & -1 \end{pmatrix} \tag{11}$$

Note here that $f'$ is positive. Simplifying the notation and letting $f'_{x_k} = f'(b \pm g_{xx} x_k^*)$

$$\det(J_e) = \frac{1}{\tau}\det\begin{pmatrix} -1 & g_{ee} f'_{e_2} \\ g_{ee} f'_{e_1} & -1 \end{pmatrix} = \frac{1}{\tau}(1 - g_{ee}^2 f'_{e_1} f'_{e_2}) = 0 \tag{12}$$

$$\det(J_i) = \frac{1}{\tau}\det\begin{pmatrix} -1 & -g_{ii} f'_{i_2} \\ -g_{ii} f'_{i_1} & -1 \end{pmatrix} = \frac{1}{\tau}(1 - g_{ii}^2 f'_{i_1} f'_{i_2}) = 0 \tag{13}$$

Thus there will be at least a saddle node bifurcation whenever $g_{ee} = \sqrt{\frac{1}{f'_{e_1} f'_{e_2}}}$ or $g_{ii} = \sqrt{\frac{1}{f'_{i_1} f'_{i_2}}}$.

Note that for feedback inhibition we cannot even have a saddle-node bifurcation. If we examine the firing-rate model

$$\tau \frac{de_1}{dt} = -e_1 + f(b - g_{ii} i_1) \tag{14}$$

$$\tau \frac{di_1}{dt} = -i_1 + f(b + g_{ee} e_1) \tag{15}$$

we see that the Jacobi an can never have a 0 eigenvalue.

$$\det(J_{ei}) = \frac{1}{\tau}\det\begin{pmatrix} -1 & -g_{ie} f'_{i_1} \\ g_{ei} f'_{e_1} & -1 \end{pmatrix} = \frac{1}{\tau}(1 + g_{ei} g_{ie} f'_{e_1} f'_{i_1}) > 0 \tag{16}$$

This is because all terms are restricted to be positive. Thus feedback inhibition can only have one stable fixed point.

The next step here to prove that we have a cusp bifurcation, and not just a saddle-node bifurcation, is to prove that the quadratic part of the system degenerates. To do this we need to find a center manifold, expand along that manifold, and show that the quadratic term can be zero. We begin by finding the center manifold for the two systems. This is a one dimensional surface such that $e_2 = M_e(e_1)$ and $i_2 = M_i(i_1)$.

We can calculate the manifold by noting that

$$\frac{de_2}{dt} = \frac{dM_e(e_1)}{dt} = \frac{dM_e}{de_1}\frac{de_1}{dt}. \tag{17}$$

and

$$\frac{di_2}{dt} = \frac{dM_i(i_1)}{dt} = \frac{dM_i}{di_1}\frac{di_1}{dt}. \tag{18}$$

Solving for $M_i$ and $M_e$ gives us

$$\frac{dM_e}{de_1} = \frac{-M_e(e_1) + f(b + g_{ee}e_1)}{-e_1 + f(b + g_{ee}M_e(e_1))} \tag{19}$$

$$\frac{dM_i}{di_1} = \frac{-M_i(i_1) + f(b - g_{ii}i_1)}{-i_1 + f(b - g_{ii}M_i(i_1))} \tag{20}$$

In this particular case the excitatory manifold is easier to algebraically determine the excitatory manifold, so we will proceed only with the math for the excitatory subsystem here. The inhibitory center manifold $M_i$ must be solved for numerically.

The excitatory manifold is solvable with the ansatz $e_2 = e_1 = M_e(e_1)$. Plugging this in gives us,

$$\frac{dM_e}{de_1} = \frac{-e_1 + f(b + g_{ee}e_1)}{-e_1 + f(b + g_{ee}e_1)} = 1 \tag{21}$$

.

$\frac{dM_e}{de_1} = 1$ can be solved very easily by separation of variables, giving us $M_e(e_1) = e_1$, thereby proving that the ansatz is valid.

Continuing onward, we have the dynamics constrained on a one dimensional manifold $e := M_e$ and $M(i) := M_i$. Here, we have switched to $e$ and $M(i)$ to emphasize the fact that $i$ is no longer in the same coordinates as $i_1$ and $i_2$.

$$\tau\frac{de}{dt} = -e + f(b + g_{ee}e) \tag{22}$$

$$\tau\frac{di}{dt} = -i + f(b - g_{ii}M(i)) \tag{23}$$

$$\tag{24}$$

We can now expand this using a Taylor series.

$$\begin{aligned}\tau\frac{de}{dt} =& - e^* + f(b + g_{ee}e^*) + (e - e^*)\left(g_{ee}f'(b + g_{ee}e^*) - 1\right) \\ &+ \frac{g_{ee}^2}{2}(e - e^*)^2 f''(b + g_{ee}e^*) + \frac{g_{ee}^3}{6}(e - e^*)^3 f^{(3)}(b + g_{ee}e^*) + O\left(e^4\right)\end{aligned}$$

$$\begin{aligned}\tau\frac{di}{dt} =& - i^* + f(b - g_{ii}M(i^*)) + (i - i^*)\left[-g_{ii}M'(i^*)f'(b - g_{ii}M(i^*)) - 1\right] \\ &+ \frac{g_{ii}^2(i - i^*)^2}{2}\left[M'(i^*)^2 f''(b - g_{ii}M(i^*)) + M''(i^*)f'(b - g_{ii}M(i^*))\right] \\ &- \frac{g_{ii}^3 i(i - i^*)^3}{6}\left[M'(i^*)^3 f^{(3)}(b - g_{ii}M(i^*)) + 3M'(i^*)M''(i^*)f''(b - g_{ii}M(i^*)) + M^{(3)}(i^*)f'(b - g_{ii}M(i \right. \\ &+ O\left(i^4\right)\end{aligned}$$

Using a change of variables, we can convert the above equations into two depressed cubics giving us

$$\tau\frac{d\hat{e}}{dt} = k_e + a_e\hat{e} - \hat{e}^3$$

$$\tau\frac{d\hat{i}}{dt} = k_i + a_i\hat{i} - \hat{i}^3$$

where

$$\hat{e} = e - \frac{f''}{(g_{ee}f^{(3)})} \tag{25}$$

$$k_e = \frac{8\left(\frac{3}{4}(f)\left(f^{(3)}\right)^2 + \frac{1}{40}\left(f''\right)^3 - \frac{27}{50}\left(f' - \frac{1}{g_{ee}}\right)\left(f''\right)\right)}{g_{ee}^3\left(f^{(3)}\right)^3}$$

$$a_e = \frac{\frac{1}{6}\left(f^{(3)}\right)\left(f' - \frac{1}{g_{ee}}\right) - \frac{1}{4}\left(f''\right)^2}{3(g_{ee}^2 f^{(3)})^2}$$

$$\hat{i} = i - \frac{1}{g_{ii}} \frac{M'(i^*)^2 f''(b - g_{ii}M(i^*)) - M''(i^*)f'(b - g_{ii}M(i^*))}{-M'(i^*)^3 f^{(3)}(b - g_{ii}M(i^*)) + 3M'(i^*)M''(i^*)f''(b - g_{ii}M(i^*)) - M^{(3)}(i^*)f'(b - g_{ii}M(i^*))}$$

$$\begin{aligned}
a_i &= \frac{1}{g_{ii}^2} A_i \alpha_i \\
\alpha_i &= \frac{(-M'(i^*)f'(b - g_{ii}M(i^*)) - 1)}{18 f(b - g_{ii}M(i^*))^2} \\
A_i &= (-M'(i^*)^3 f^{(3)}(b - g_{ii}M(i^*)) + 3M'(i^*)M''(i^*)f''(b - g_{ii}M(i^*)) \\
&\quad - M^{(3)}(i^*)f'(b - g_{ii}M(i^*)) - \frac{1}{4}(M'(i^*)^2 f''(b - g_{ii}M(i^*)) - M''(i^*)f'(b - g_{ii}M(i^*)))^2 \\
k_i &= \frac{N_i}{g_{ii}^3 d_i} \\
N_i &= 8\frac{1}{40}(M'(i^*)^2 f''(b - g_{ii}M(i^*)) - M''(i^*)f'(b - g_{ii}M(i^*)))^3 \\
&\quad - \frac{27}{50}(-M'(i^*)f'(b - g_{ii}M(i^*)) - \frac{1}{g_{ii}})(M'(i^*)^2 f''(b - g_{ii}M(i^*)) - M''(i^*)f'(b - g_{ii}M(i^*))) \\
&\quad + \frac{3}{4} f(b - g_{ii}M(i^*))(-M'(i^*)^3 f^{(3)}(b - g_{ii}M(i^*)) + 3M'(i^*)M''(i^*)f''(b - g_{ii}M(i^*)) - M^{(3)}(i^*)f'(b - \text{[illegible]} \\
d_i &= (-M'(i^*)^3 f^{(3)}(b - g_{ii}M(i^*)) + 3M'(i^*)M''(i^*)f''(b - g_{ii}M(i^*)) - M^{(3)}(i^*)f'(b - g_{ii}M(i^*)))^3
\end{aligned} \tag{26}$$

Whenever $k_e$, and $a_e$, or $k_i$, and $a_i$ are $0$, then the two systems undergo a cusp bifurcation. While the exact values of the cusp bifurcation is complicated, especially in the case of the inhibitory system, it still can undergo the bifurcation.

### Coupling mutual excitation with inhibition

Now that we have calculated the normal form for both of subsystems, we can couple them together. We don't need to explicitly use the normal form at first. Rather its mere existence will be used later. We begin by introducing the coupling terms $\epsilon$. The corresponding equation becomes

$$\tau\frac{de_1}{dt} = -e_1 + f(b + g_{ee}e_2 - \epsilon[g_{ie}\theta_{i11}i_1 + g_{ie}\theta_{i12}i_2]) \tag{27}$$
$$\tau\frac{de_2}{dt} = -e_2 + f(b + g_{ee}e_1 - \epsilon[g_{ie}\theta_{i21}i_1 + g_{ie}\theta_{i22}i_2]) \tag{28}$$
$$\tau\frac{di_1}{dt} = -i_1 + f(b - g_{ii}i_2 + \epsilon[g_{ei}\theta_{e11}e_1 + g_{ei}\theta_{e12}e_2]) \tag{29}$$
$$\tau\frac{di_2}{dt} = -i_2 + f(b - g_{ii}i_1 + \epsilon[g_{ei}\theta_{e21}e_1 + g_{ei}\theta_{e22}e_2]) \tag{30}$$

Here, $\theta_{k1}^2 + \theta_{k2}^2 = 1$ and $\theta_{kx} > 0$. This allows the coupling between the connections to be arbitrary. We can think of $\theta$ as a way to parameterize the asymmetries in synaptic weights between the coupling.

Next, by expanding the function $f$ as a Taylor series with respect to $\epsilon$ centered at $\epsilon = 0$, and then disregarding order $\epsilon^2$ and above, we get

$$\tau\frac{de_1}{dt} = -e_1 + f(b_e + g_{ee}e_2) - \epsilon f'(b_e + g_{ee}e_2)(g_{ie}\theta_{i11}i_1 + g_{ie}\theta_{i12}i_2) \tag{31}$$
$$\tau\frac{de_2}{dt} = -e_2 + f(b_e + g_{ee}e_1) - \epsilon f'(b_e + g_{ee}e_1)(g_{ie}\theta_{i21}i_1 + g_{ie}\theta_{i22}i_2) \tag{32}$$
$$\tau\frac{di_1}{dt} = -i_1 + f(b_i - g_{ii}i_2) + \epsilon f'(b_i - g_{ii}i_2)(g_{ei}\theta_{e11}e_1 + g_{ei}\theta_{e12}e_2) \tag{33}$$
$$\tau\frac{di_2}{dt} = -i_2 + f(b_i - g_{ii}i_1) + \epsilon f'(b_i - g_{ii}i_1)(g_{ei}\theta_{e21}e_1 + g_{ei}\theta_{e22}e_2) \tag{34}$$
$$\tag{35}$$

We can then make the substitution for the stable attractive manifolds we made above such that $e = e_2 = M(e_1) = e_1$ and $i = i_2 = M_i(i_1)$.

$$\tau\frac{de}{dt} = -e + f(b_e + g_{ee}e) - \epsilon f'(b_e + g_{ee}e)(g_{ie}(\theta_{i11} + \theta_{i21})i + g_{ie}(\theta_{i12} + \theta_{i22})M_i(i)) \tag{36}$$

$$\tau\frac{di}{dt} = -i + f(b_i - g_{ii}M_i(i)) + \epsilon f'(b_i - g_{ii}M_i(i))(g_{ei}(\theta_{e11} + \theta_{e21})e + g_{ei}(\theta_{e12} + \theta_{e22})e) \tag{37}$$

For ease of notation let $\theta_{i_1} = \theta_{i11} + \theta_{i21}$ and $\theta_{i_2} = \theta_{i12} + \theta_{i22}$, and $\theta_{e_1} = \theta_{e11} + \theta_{e21}$ and $\theta_{e_2} = \theta_{e12} + \theta_{e22}$ We can now expand the equation in terms of $e$ and $i$ giving us

$$\begin{aligned}\tau\frac{de}{dt} =& - e^* + f(b + g_{ee}e^*) + (g_{ee}f'(b + g_{ee}e^*) - 1) + \frac{g_{ee}^2(e-e^*)^2}{2}f''(b + g_{ee}e^*) \qquad (38)\\ &+ \frac{g_{ee}^3(e-e^*)^3}{6}f^{(3)}(b + g_{ee}e^*) + O\left(e^4\right) - \epsilon f'(b_e + g_{ee}e)(g_{ie}\theta_{i1} + g_{ie}\theta_{i_2}M_i'(i))(i - i^*) + O(i^2)\end{aligned}$$

$$\begin{aligned}\tau\frac{di}{dt} =& - i^* + f(b - g_{ii}M(i^*)) + (i - i^*)\left[-g_{ii}M'(i^*)f'(b - g_{ii}M(i^*)) - 1\right] \qquad (39)\\ &+ \frac{g_{ii}^2(i-i^*)^2}{2}\left[M'(i^*)^2f''(b - g_{ii}M(i^*)) + M''(i^*)f'(b - g_{ii}M(i^*))\right] \qquad (40)\\ &- \frac{g_{ii}^3(i-i^*)^3}{6}\left[M'(i^*)^3f^{(3)}(b - g_{ii}M(i^*)) + 3M'(i^*)M''(i^*)f''(b - g_{ii}M(i^*)) + M^{(3)}(i^*)f'(b - g_{ii}M(i^*\right.\\ &+ O\left(i^4\right) + \epsilon f'(b_i - g_{ii}M_i(i))(g_{ei}\theta_{e1} + g_{ei}\theta_{e_2})(e - e^*) + O(e^2)\end{aligned}$$

Recall we can create a depressed cubic in the uncoupled system using equations 25 and 26 to get

$$\tau\frac{dE}{dt} = k_e(b, g_e e) - a_e(b, g_e e)E - E^3 - (\epsilon f'(b_e + g_{ee}e^*)(g_{ie}\theta_{i1} + g_{ie}\theta_{i_2}M_i'(i^*))(I + I^*) \tag{41}$$

$$\tau\frac{dI}{dt} = k_i(b, g_e e) - a_i(b, g_e e)I - I^3 + \epsilon f'(b_i - g_{ii}M_i(i^*))(g_{ei}\theta_{e1} + g_{ei}\theta_{e_2})(E + \frac{f''}{(g_{ee}f^{(3)})}) \tag{42}$$

where $I^* = \frac{1}{g_{ii}}\frac{M'(i^*)^2f''(b-g_{ii}M(i^*)) - M''(i^*)f'(b-g_{ii}M(i^*))}{-M'(i^*)^3f^{(3)}(b-g_{ii}M(i^*)) + 3M'(i^*)M''(i^*)f''(b-g_{ii}M(i^*)) - M^{(3)}(i^*)f'(b-g_{ii}M(i^*))}$ from equation 26. To clean everything up a bit, we can recast our equations as

$$\tau \frac{dE}{dt} = k_e^* - a_e E + E^3 - \epsilon a_{ie} I \tag{43}$$

$$\tau \frac{dI}{dt} = k_i^* - a_i I - I^3 - \epsilon a_{ei} E \tag{44}$$

where $a_{ie} = g_{ie}\theta_{i1} + g_{ie}\theta_{i2}M'$, $a_{ei} = g_{ei}\theta_{e1} + g_{ei}\theta_{e2}$, $k_e^* = k_e + \frac{f''}{\left(g_{ee}f^{(3)}\right)}$, and $k_i^* = k_i + I^*$.

### Supplementary Figures

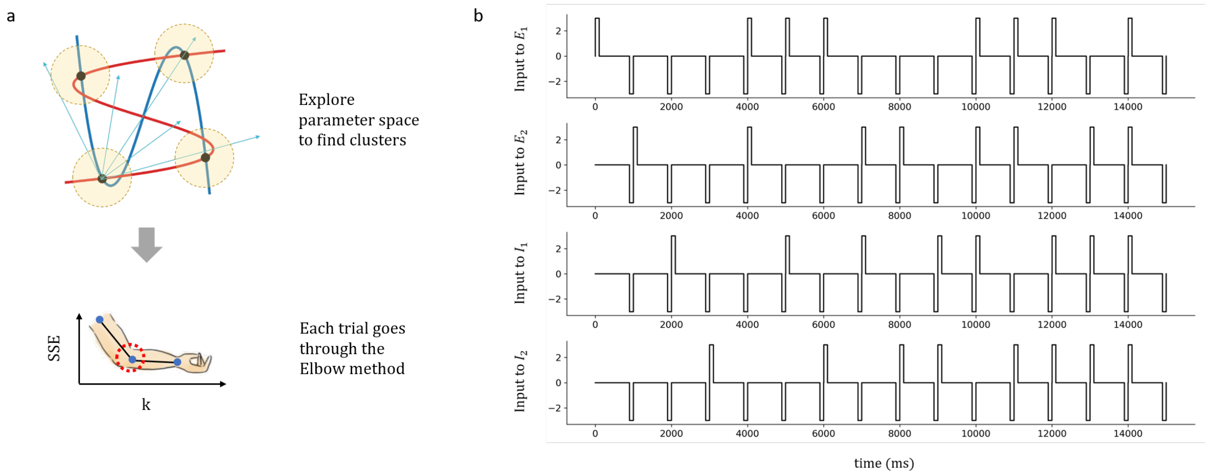


Figure S1: Protocol for statistical analysis of equilibrium points. (a) *Top:* Schematic diagram of attractors in a motif. Each attractor has its own basin of attraction, so that once the system is pushed near the attractor, it stays near the attractor. The protocol is designed to explore different state spaces so as to find as many attractors as possible. *Bottom:* The method we used to determine the number of equilibrium points is the elbow method. Specifically, when the algorithm hits the number of correct clusters $k$ in the data, it will result in an elbow-like dip in $SSE$, as shown in the graph. (b) The current stimulation given to the four neurons in one trial. The duration for the positive pulse and reset pulses are both 100 ms, while the interval in between the pulses are 800-900 (ms) – the precise interval value doesn't matter, as long as they are longer than the transient activities.

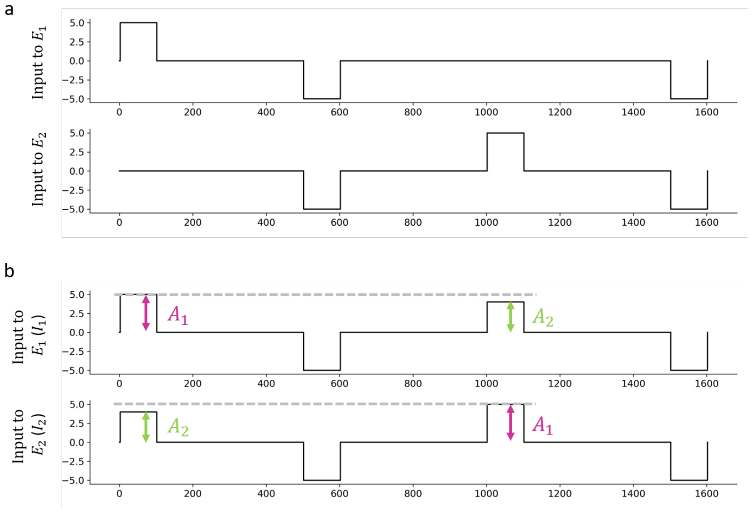


Figure S2: Input protocol for decision making and switch. (a) The input protocol for switch. The duration of the pulses are 100 ms, and the interval in between is 400 ms. The strength of the positive and reset pulses is 5 and -5 nA, respectively. (b) The input protocol for decision making. Note that $A_1$ and $A_2$ have different amplitudes ($A_1$ =5 nA and $A_2$ =4 nA), representing how the input signal strength is different for the two neurons. The reset pulse is -5 nA. No noise is included in this simulation, but including noise does not change the results qualitatively.

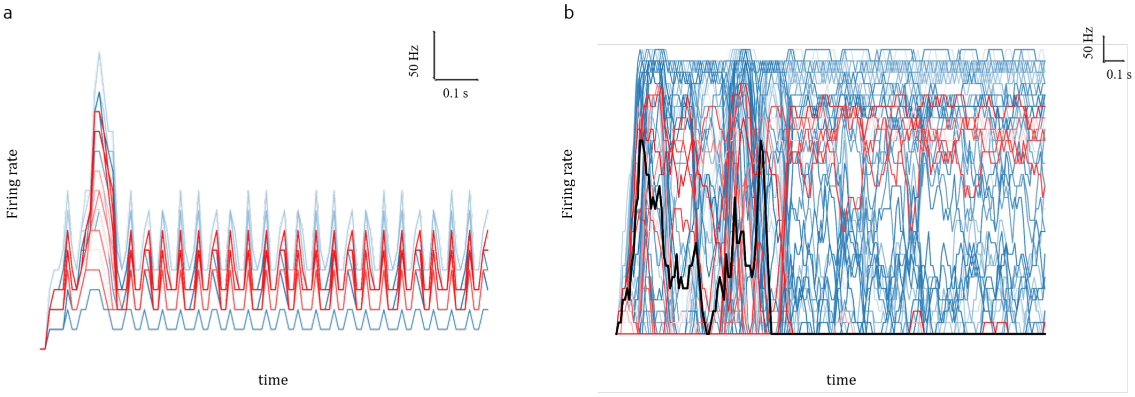


Figure S3: CPG and toggle in large random network. (a) Firing rate of neurons in the network performing CPG, blue is excitatory and red is inhibitory. (b) Firing rate of neurons in the network performing toggle, blue is excitatory and red is inhibitory. Black is one particular neuron used to showcase toggle more clearly. The parameters for toggle is: $g_{ee} = 25$ $(\mu S)$, $g_{ei} = 15$ $(\mu S)$, $g_{ie} = 150$ $(\mu S)$, $g_{ii} = 50$ $(\mu S)$, $b_E = 2.5$ $(nA)$, $b_I = 0$ $(nA)$. The parameters for CPG is: $g_{ee} = 25$ $(\mu S)$, $g_{ei} = 25$ $(\mu S)$, $g_{ie} = 50$ $(\mu S)$, $g_{ii} = 25$ $(\mu S)$, $b_E = 1$ $(nA)$, $b_I = 0$ $(nA)$.

### Supplementary Tables

#### Supplementary Table 1

| Parameter Space for Equilibrium Points Test | | | |
|---|---|---|---|
| Parameter | Value | Parameter | Value |
| $g_{ee}$ $(\mu S)$ | 90, 105, 120, 135, 150 | Amplitude (nA) | 1, 3, 5, 7 |
| $g_{ii}$ $(\mu S)$ | 30, 40, 50, 60, 70, 80, 90 | Combination | $C_1^4$, $C_2^4$, $C_3^4$, $C_4^4$ |
| $g_{ie}$ $(\mu S)$ | 15, 30, 45 | Bias (nA) | 0, 0.55, 1.55 |

Table S1: Parameter space for determining the equilibrium points of the motifs. $g_{ei}$ is fixed at 30 $\mu S$. Amplitude represents the amplitude of the positive pulses given in units of nA. The amplitude for the reset pulses are fixed at -3 nA. Combination reflects the different combinations of neurons that are given the pulse. For example, at a given time period, neurons $E_1$ and $E_2$ might receive current pulses while neurons $I_1$ and $I_2$ do not. This is iterated through all different combinations. Bias represents the underlying bias currents that all neurons receive.

#### Supplementary Table 2

| Parameter Space for CPG Test | |
|---|---|
| Parameter | Value |
| $g_{ei}$ $(\mu S)$ | 0 to 60, increments of 5 |
| $g_{ie}$ $(\mu S)$ | 0 to 200, increments of 10 |

Table S2: Parameter space for determining the CPG of the motifs. $g_{ee}$ is fixed at 65, $g_{ii}$ at -5, the bias current for excitatory neurons is $b_E = 0.5$, and the bias current for inhibitory neurons is $b_I = -0.5$.

#### Supplementary Table 3

| CRIREL function parameters | | | | | | |
|---|---|---|---|---|---|---|
| Function | $g_{ee}$ $(\mu S)$ | $g_{ei}$ $(\mu S)$ | $g_{ie}$ $(\mu S)$ | $g_{ii}$ $(\mu S)$ | $b_E$ $(nA)$ | $b_I$ $(nA)$ |
| Switch | 110 | 0 | 0 | 0 | 0 | 0 |
| Toggle/sCPG | 65 | 30 | -200 | -5 | 0.5 | -0.5 |
| Decision | 100 | 70 | -70 | -70 | 2.4 | 0 |
| Toggle-decide | 10 | 70 | -70 | -25 | 2.4 | 0 |
| aCPG | 10 | 70 | -70 | -25 | 2.4 | 0 |
| Memory | 75 | 45 | -2 | -30 | 0.5 | 0.5 |
| AND | 30 | 30 | -60 | -30 | -0.2 | -0.6 |
| XOR | 30 | 30 | -60 | -30 | 0.3 | 0.55 |

Table S3: The parameters used to produce the CRIREL circuit functions in the figures. $b_E$ is the bias current for excitatory neurons, and $b_I$ is the bias current for inhibitiory neurons. The parameters for the neuronal and synaptic model follows the parameters given in the methods section unless otherwise specified. The parameters for amplitude-based decision is more complicated, and is listed here instead: $g_{ee} = 50$ $(\mu S)$, $g_{ie}$=-4000 $(\mu S)$, $g_{E_1 to I_2}$=25 $(\mu S)$, $g_{E_1 to I_1}$=20 $(\mu S)$, $g_{I_1 to I_2}$=70 $(\mu S)$, $g_{I_2 to I_1}$=30 $(\mu S)$. Also, the membrane time constant $\tau$ is 80 (ms) and capacitance $C$ is 1 $nF$ for $I_2$.

#### Supplementary Table 4

| Large Network Parameters | | | |
|---|---|---|---|
| Parameter | Value | Parameter | Value |
| $p_{ee}$ | 0.5 | $g_{ee}$ $(\mu S)$ | 20 |
| $p_{ei}$ | 0.5 | $g_{ei}$ $(\mu S)$ | 6 |
| $p_{ie}$ | 0.5 | $g_{ie}$ $(\mu S)$ | 0 to 28, increments of 2 |
| $p_{ii}$ | 0.5 | $g_{ii}$ $(\mu S)$ | 0 to 29, increments of 1 |

Table S4: Connection probability and synaptic weight of large network. $p_{ij}$ represents the connection probability of a neuron of type $j$ to type $i$, where $i, j \in E, I$. $g_{ij}$ represents the synaptic weight of a neuron of type $j$ to type $i$.
